# Untargeted Metabolomics of Plant Samples using HPLC-DAD and Gaussian Mixture Models

**DOI:** 10.1101/2025.11.10.687686

**Authors:** Justin Gambill, Chase Mason, Jordan A. Dowell

## Abstract

**Premise:** Plants produce millions of different chemical compounds, contributing greatly to their physiology and evolutionary trajectories. Most untargeted metabolomic methods are inaccessible, either due to upfront instrument costs or intensive technical training. More accessible methods using diode array detectors often only utilize a few wavelengths, preventing high-throughput observation of total metabolic diversity.

**Methods:** Leaves from the genera *Betula*, *Magnolia, Rosa,* and *Viburnum* were collected, dried and ground, extracted, and analyzed by HPLC-DAD. Chromatographic data was then processed in a curated R pipeline, and resulting resolved peaks were clustered by absorbance spectra using Gaussian Finite Mixture Models (GMMs). To assess clustering, GMM was compared to a more traditional linear discriminant analysis (LDA) method, with clusters identified through literature searches.

**Results:** Significant associations between the abundances of chemical classes and whole-metabolome alpha and beta diversity indices were recovered. In general, GMMs performed better than other classification methods like LDA, especially between classes that share common features like non-flavonoid phenolics and flavonoids.

**Discussion:** We show that our method can easily extract relevant class-level diversity of metabolite profiles among closely related species, genotypes, and ecotypes. Regardless of underlying research question, our method extends the usage of DAD beyond restricted targeted analyses and increases the accessibility of untargeted metabolomics.

## Introduction

Plants provide a unique challenge to metabolomics research. The metabolome of a typical plant can contain several orders of magnitude more metabolites than what can be found in a typical animal species, in addition to a much higher variability between the chemical profiles of differing species – even within the same genus (Fiehn, 2002; Domingo-Fernandez et al., 2023). Untargeted metabolomics, which aims to separate as many compounds as possible from a single sample, provides a powerful workflow for observing variation in chemical diversity across entire groups of plants.

Metabolomics traditionally employs two steps: a separation step to isolate as many chemical compounds in a biological sample as possible, usually using a gas or liquid chromatograph; and an identification step, usually with a mass spectrometer, which identifies compounds based on ion fragmentation patterns (i.e., mass to charge ratios; Kim et al., 2012). Mass spectrometry, although powerful, presents significant accessibility barriers, both in the cost of the instrument (upwards of $100,000 USD) and the time required to operate it efficiently and safely. Therefore, providing methods to enhance the insights available from other detectors will increase the accessibility of phytochemistry research to laboratories without reliable access to mass spectrometers, as well as enable lower-cost, high-throughput applications for all researchers.

### Diode Array Detection Analysis of Untargeted Metabolomic Data

Diode array detectors (DADs) measure the absorbance of chemical compounds across a range of wavelengths of light, typically from the far ultraviolet (UV) range (190nm) to the near-infrared range (900nm). Absorbance across the range can be measured simultaneously, allowing for multiple features to be used for identification. Often, these detectors are coupled with high-performance liquid chromatography or capillary electrophoresis, allowing for the non-destructive sampling and analysis of a broad range of compounds in complex mixtures. Non-destructively, in this case, means that compounds separated and identified via HPLC-DAD can be fractionated, collected, and used in follow-up mechanistic studies (e.g., toxicology, bioprospecting, isolation for pollinator attraction studies). Unlike mass spectrometry, however, DADs cannot provide extensive structural information beyond the portion of the chemical compounds’ structure that interacts with light for each compound, or “peak”, as separated by high-performance liquid chromatography (HPLC) (Figure 1B). However, the absorbance spectrum of compounds detected by the DAD is indicative of common structural elements of specific chemical classes that interact with light (chromophores), often minimally affected by variation in functional group addition (Taniguchi et al., 2023a). Therefore, DAD data provide a cost-effective (approximately $20,000 USD) and robust method for obtaining class-specific metabolomic data of plant samples.

**Figure 1.**
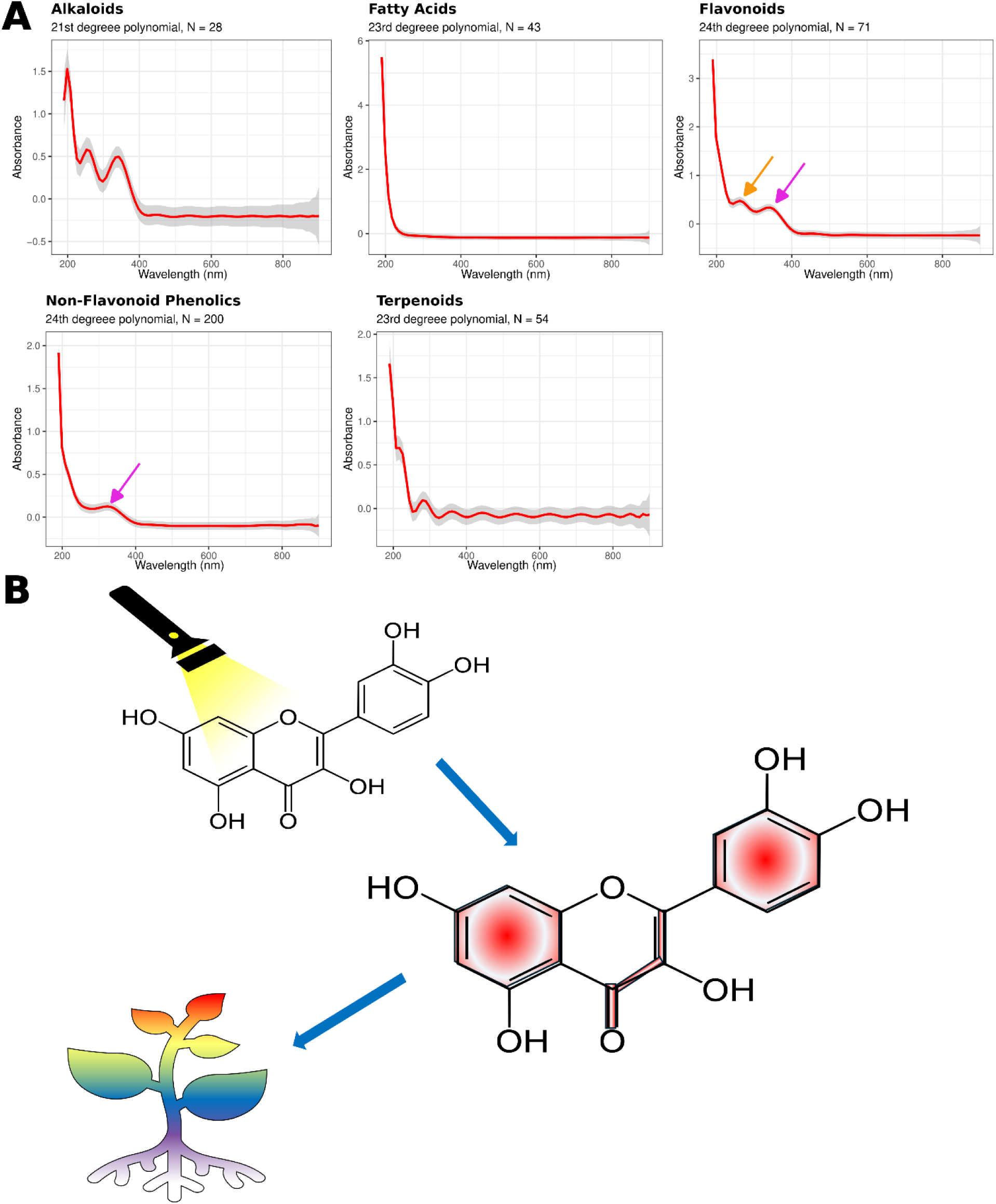
Chromophore classes described by Gaussian Mixture Model classification. A) Reconstructed absorbance spectra generated after GMM clustering and polynomial regression analysis. Regression lines are depicted in red with 97% confidence interval, shaded in grey. N equals the number of unique peaks belonging to each class. Wavelength is measured in nanometers. Magenta arrows indicate common sources of classification confusion between classes that share similar chromophore structures. Orange arrows show small differences that the GMM can use to distinguish between the similarities described by the magenta arrows. B) Schematic diagram of chromophores interacting with light. A light source will excite conjugated pi-bond systems, like those found in quercetin (chemical structure shown). The excitation will force electrons into a higher energy orbital, after which their return to their “ground state” its color. Chromophore is outlined in red to show photoactive portions of the molecule. Multicolor gradient of the plant is shown to illustrate the myriad of unique photoreactive properties of each individual chromophore found within any given sample.

### Chromophores and Gaussian Finite Mixture Modeling

Chromophores are the central structures of compounds that interact with light, giving a particular compound or class of similarly structured compounds (e.g., terpenoids, flavonoids, and alkaloids) a characteristic absorption spectrum that can be useful for classification purposes.

Chromophores primarily form from conjugated pi-bond systems, where non-bonding electrons are excited when introduced to light. This excitation forces a molecular electron to shift from a high-energy molecular orbital to an empty low-energy molecular orbital, contributing to a measurable absorbance value indicative of the chromophore’s structure, where changes to the conjugated pi-bond system alter the absorbance pattern (Figure 1). While many traditional UV-Vis spectroscopy methods may only look at the maximal absorbance of a few specific wavelengths, incorporating information from the entire absorbance spectrum may lead to better classification or delineation among chemical classes with subtle differences in their chromophores. In practice, DADs attached to HPLC systems typically measure absorbance from 190 to 900 nm, yielding a wide range of potentially detectable chromophores. However, the variation among chromophores and the need to assign individual compounds to chemical classes necessitate clustering methods. To our knowledge, no method exists that attempts to cluster peaks based on their entire absorption spectra, most likely reflecting the primary goal of accurately quantifying a small number of compounds rather than obtaining a broad characterization of relative chemical diversity. However, we hypothesize that clustering algorithms that maximize weights given to common features are well-suited to describing relative class-level chemical diversity, as absorbance similarities among compounds that share a common chromophore reflect fundamental physiochemical interactions. Additionally, as each chemical class has shared structural features (e.g., the unique chromophore), one can consider the absorbance pattern of each class as a probability distribution, where variance can be attributed to functional group and side chain additions to the common structure.

Gaussian Finite Mixture Modeling (GMM) is an unsupervised clustering algorithm that groups components based on the probability that each component belongs to a particular cluster. GMMs fit clusters based on the shape and distribution of the data, making them powerful tools for clustering chemical spectra based on similar features. GMMs combine estimation-maximization algorithms with model-based hierarchical clustering (Scrucca et al., 2016). The model assumes that all data points can be modeled as a family of Gaussian distributions, through which an optimized local maximum can describe data points with similar eigen structures. The generalized model is given by Equation 1,

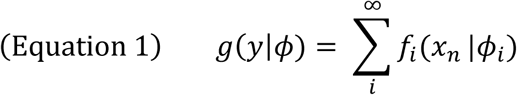

where **x** represents the eigenvalues of each sample, ϕ is a vector that describes the underlying probability distribution variables (mean and covariance matrix structure that describe distribution size and shape), and y corresponds to a hyperparameter that defines the local maximum of the Gaussian of each individual cluster in the dataset. The model we propose to estimate chemical diversity uses the expectation maximization algorithm described by Hasselblad (1966) that assumes that each peak in the chromatogram is expected to cluster around a designated number of local maxima (y), followed by a maximum-likelihood optimization step that fits parameters (ϕ) given the sample’s underlying eigenstructure to a given local maximum of the integrated model. In this case, each local maxima of the function corresponds to a particular chromophore, and the covariance is derived from functional group and side chain additions to the main chromophore. The model is then iteratively run with an increasing number of expected clusters until the optimal number of clusters is achieved based on minimizing the Bayesian information criterion.

Once clustering is complete, samples within each cluster are scaled and combined into a single data set, where a polynomial regression is fit to reconstruct the average absorbance spectra. The reconstructed spectra enable class distinction through comparison with library standards in databases such as Reaxys and other published spectra in the literature (Haghi & Hatami, 2010; Hisiger & Jolicoeur, 2007; Taniguchi et al., 2023a; Walker & Hawkins, 1952). This classification can then be used as a phenotypic marker beyond peak number and abundance, further deepening our understanding of plant chemical diversity.

The objective of this study is to develop a method that can extract chemical class information from high-throughput, untargeted HPLC-DAD data. We first optimize an analytical chemistry method for four taxonomically and chemically distinct genera: *Betula, Magnolia, Rosa,* and *Viburnum*. We then employ two clustering approaches to describe the chemical diversity extracted from each sample. Our proposed GMM model was first applied to describe the data set initially. GMM classifications were then used as predictor values for a subsequent principal component analysis coupled with a linear discriminant analysis. The resulting GMM classes were then used to test whether or not a derived chemical diversity index (e.g., the Shannon Diversity of the metabolome) could explain the variation in class abundances across each genus.

## Materials and Methods

### Plant Tissue Collection

Four plant genera were included in this study – *Betula, Magnolia, Rosa,* and *Viburnum* – to provide a vast array of chemical diversity. Specimens of each species were sampled from within the living collections of the Arnold Arboretum of Harvard University (Boston, Massachusetts) during the summer of 2016, including 27 species of *Betula* (58 specimens), 18 species of *Magnolia* (57 specimens), 57 species of *Rosa* (87 specimens), and 41 species of *Viburnum* (103 specimens), including a range of subspecies and taxonomic varieties of species within each genus (Appendix 1). For all members of each genus, leaf sampling was performed over one or two days within the same week to ensure similar leaf ontogeny and environmental conditions in the common garden conditions of the Arnold Arboretum. Undamaged green leaves were removed from each specimen and placed in zip-sealed plastic bags, which were kept in an ice-cooled chest until they were returned to the laboratory. The leaves were then dried in a forced-air drying oven at 60°C for 3 days until they reached a constant weight. Dried leaves were ground into a fine homogenized powder for chemical analysis.

### Plant Material Extraction

Each plant sample was extracted first in 800 µL of a 1:1 chloroform:methanol solution, shaken to homogenize, and sonicated on ice for 30 minutes. For initial phase separation, 400 µL of HPLC-grade water was added. Samples were then centrifuged for 10 minutes at 1000 rpm at 4°C using an Eppendorf Centrifuge 5810 R (Hamburg, Germany). 300 µL of the upper aqueous phase was aliquoted into a new 96-well plate and centrifuged again at 1000 rpm at 4°C for 5 minutes.

Finally, a 200 µL aliquot was filtered through a 0.45 µm hydrophilic, low protein-binding Durapore membrane filter (VWR Scientific, Randor, PA), then sealed for analysis. All solvents were purchased through VWR (Randor, PA).

### Chromatographic Separation

All samples were run on an Agilent 1260 Infinity II High-Performance Liquid Chromatography (HPLC) system (Santa Clara, CA) equipped with a multisampler and quaternary pump. The chromatography parameters were optimized for each genus separately, based on the peak diversity of pooled samples from species within each genus (Appendix 1). Data was blank-subtracted and smoothed using a Gaussian smoothing algorithm in Agilent CDS-OpenLabs software (Santa Clara, CA), and then downloaded for further processing.

### Peak Alignment

Peak alignment was achieved using a modified workflow with the chromatographR package in R version 4.4.1 (Bass, 2023). Samples were initially pre-processed and baseline-corrected by redesignating the time window to exclude the column cleaning and solvent re-equilibration steps of the chromatographic methods using the *preprocess* function. Peak alignment and smoothing were performed using a parametric time-warping algorithm across the entire wavelength range (190-900nm). Peak-specific absorbance spectra were then extracted from the *warp* object and used for clustering analysis. Peak areas were then integrated using an exponential-Gaussian hybrid function, and homologous peaks were combined based on maximum spectral similarity by calculating Pearson’s correlations across all samples and combining peaks with a correlation above 0.9 and a retention time difference smaller than 0.05 minutes. In many cases, peaks at the end of the chromatogram represent fragments of the column or are not measurable due to column bleed. This results in the algorithm assigning multiple features to the same retention time, differing only by the maximum absorbance used for quantification. For example, seven peaks could be identified at a retention time of 6 minutes (given the overall runtime of the method was 6.5 minutes), with each peak being attached to a quantification wavelength of 300, 305, 310, 400, and 500 nm. As DAD lacks true structural information, we are unable to discern whether each peak represents either a unique compound or multiple measurements of the same compound. Therefore, these peaks were considered erroneous and were removed to ensure that the peaks retained in the data set are those with unique absorption spectra, which are used in the clustering analysis. The resulting dataset describes all unique peaks across a set of samples. Peak areas were then normalized by the sample weight and multiplied by a concentration factor, which was the ratio between the aliquot and injection volumes.

### Peak Clustering and Spectra Reconstruction

For homologous peaks across samples, we selected the peak from the sample with the highest absorbance to build a representative library of chemical diversity within the study. We chose the peak with the highest absorbance, as the Beer-Lambert law states that the change in the quantity of a specific compound between two samples is equivalent to the change in absorbance.

Therefore, the peak with the highest absorbance has the highest concentration of a specific compound among samples, leading to greater confidence in the accuracy of the compound’s absorbance spectrum (as a higher signal-to-noise ratio is observed for a compound with higher concentration). The resulting library was then transformed so that each column corresponded to a different wavelength, and each row represented a distinct peak. The resulting data set was then clustered by the following pipeline using functions available in the *mclust* and *poly* packages in R. Initial model parameters were first designated by the maximum BIC using the *mclustBIC()* function. The resulting vector of model coefficients was then used to optimize a Gaussian mixture model using the *mclust()* function. Cluster assignments were then extracted and used for further spectral analysis.

To control potential overfitting of the GMM spectral reconstruction of each cluster, a polynomial regression was fitted with a degree selected based on the lowest root mean squared error. The first complete spectrum (in our case, wavelengths ranging from 190 to 900nm, in increments of 5 nm). The final model was optimized following a 10-fold cross-validation using functions available in the *caret* package in R. Resulting spectra were then further described by a chemical class identifier (i.e., flavonoids) based on available spectra in the Reaxys database and literature (Haghi & Hatami, 2010; Hisiger & Jolicoeur, 2007; Taniguchi et al., 2023a; Walker & Hawkins, 1952).

To compare our clustering approach with more traditional methods, the classification results from the GMM were used as predictor labels for further model testing. The resulting chemical classes were then compared to a clustering analysis that applied PCA via singular value decomposition to the correlation matrix of the dataset coupled to linear discriminant analysis (LDA), with the number of clusters defined by the number of chemical classes extracted from the Gaussian mixture modeling approach. PCA, coupled with LDA, was employed as an example of more traditional chemometric approaches (Kharazian et al., 2024; Sammarco et al., 2023). LDA was applied to the values of the first four principal components, encompassing >90% of the observed variation in the data set, to compare the classification strength against GMM clustering. PCA was then used for visualizing the resulting chemospace. Labels derived from GMM analysis were used as predictors, and subsequent confusion matrices were analyzed to examine the differences in classification abilities between the two models. PCA-LDA model performance was assessed through a confusion matrix, Cohen’s Kappa, and overall model accuracy, sensitivity, and selectivity. Cohen’s Kappa measures the level of agreement between the observed classifications made by the PCA-LDA model, against the classifications made by the GMM. Cohen’s Kappa is calculated as the difference between the observed agreement and the hypothetical agreement, divided by one minus the hypothetical agreement. Values closer to 1 indicate stronger agreement between the observed classifications and the hypothetical classifications. Specificity and sensitivity were also calculated following PCA-LDA. Specificity measures the model’s ability to accurately identify true negative classifications, and sensitivity measures the model’s ability to accurately identify true positive classifications. The resulting R code for peak alignment, processing, and clustering can be found at the following link: https://github.com/justingambill/ChemicalClusteringbyGMM.

### Phylo-comparative Analyses

Resulting metabolic profiles and chemical classes were used to perform phylogenetic generalized least squares (PGLS) regressions to test whether or not the derived chemical diversity could explain variation in chemical class abundance across a genus. Phylogenies for the genera *Betula, Magnolia, Rosa,* and *Viburnum* were constructed using freely and publicly available gene sequences from NCBI (Appendix 2). A modified pipeline using *phylotaR,* NCBI BLAST search (Bennet et al., 2018; Sanderson et al., 2008; Wang et al., 2025), and MAFFT (Katoh et al., 2013) with default parameters was used to align and cluster similar gene sequences for ∼4 genes per genus into a supermatrix. This supermatrix file was used to produce an initial maximum likelihood phylogeny using RAxML version 8 (Stamatakis, 2014). The resulting phylogenies were compared with and confirmed by the most recently published phylogenies for each genus (Fougère-Danezan et al., 2015; Schenk et al., 2008; Wang et al., 2020; Winkworth & Donoghue, 2005) and modified through pruning to include only species represented in our data sets, and re-rooting the tree to obtain a topology consistent with the most recently published phylogenies, which used multiple outgroups using the *ape* package in R (Paradis, 2019). A relaxed molecular clock was then used to convert each phylogeny into an ultrametric tree through the chronos function available in the *phytools* package in R (Revell, 2024). The model was specified as a relaxed molecular clock, the minimum age of the root and the lambda smoothing parameters were set to 1.

Chemical diversity indices were calculated using three complementary metrics: the Jaccard Similarity Index, Shannon Diversity Index, and the Inverse Simpson Index. The Jaccard Similarity Index (JSI) is calculated by measuring the ratio of the number of peaks shared by two samples over the total number of combined unique peaks within the two species. This results in a distance-similarity matrix (species by species). For Species A, we take the pairwise JSI values for Species A compared with Species B to Species N, and average the resulting vector, providing an overall metric of pairwise similarity of Species A to the rest of the genus. Calculations for the JSI were performed using a manually written function in R (version 4.4.1). The Shannon Diversity Index (SDI) measures both the abundance and diversity of a set of peaks within a given sample. Higher values indicate that the profile is more diverse and abundances are more evenly distributed across the profile; lower values indicate a less diverse chemical profile, with a few compounds existing in high abundance. The Inverse Simpson Index (ISI) is positively correlated with the SDI and measures dominance in the profile. Higher values of the ISI indicate the profile is more evenly spread, with a low probability that a random draw from the set of peaks within a sample would pull the same peak twice; therefore, lower values indicate the profile is dominated by a few peaks, with a random draw likely to pull the same few peaks more than others. The SDI and ISI were calculated using the R package *vegan* (Oksanen et al., 2025).

Our method lacks internal standards and only reports relative quantification; therefore, feature scaling was applied to reduce detection bias within each peak across samples, similar to the mass spectrometry peak processing method outlined by Bassi et al. (2025). The minimum intensity of each peak across all samples was subtracted from the maximum intensity and divided by the range of intensities. All “min-max” scaled peaks within a chemical class were then summed together within a given sample, resulting in a value that describes the total increase or decrease of a given chemical class, hereafter referred to as class abundance (Bassi et al., 2025).

Phylogenetic generalized least squares (PGLS) regressions were first performed with chemical diversity index as the predictor variable and the summed class abundance of each chemical class as the response variable, assuming a Brownian Motion model of evolution. Subsequent PGLS models were built to assess pairwise correlations between the summed class abundances. The resulting model coefficients were extracted along with their respective standard errors. PGLS regressions were accomplished using the R packages *nlme* (v3.1-152; Pinherio et al., 2021) and *phytools* (Revell, 2024).

## Results

An initial dataset was constructed from the described peaks of *Betula* (58 samples), *Magnolia* (57 samples), *Rosa* (87 samples), and *Viburnum* (103 samples). First, a “global” data set was constructed that included peaks and their associated absorbance spectra across all genera included in this study. The global dataset was clustered using both PCA-LDA and a fitted GMM. In general, GMM clustering was able to differentiate more spectral features of varying class sizes than PCA-LDA (Figure 2B and 2C). Clusters show reconstructed absorbance spectra with low standard deviations (Figure 1A). Across the four genera, five distinct chemical classes were repeatedly described: alkaloids, fatty acids, flavonoids, non-flavonoid phenolics, and terpenoids. Literature searches of published absorbance spectra assisted in structural classification of each chromophore (Haghi & Hatami, 2010; Hisiger & Jolicoeur, 2007; Taniguchi et al., 2023a; Walker & Hawkins, 1952). Genera-specific variation was captured, both as variation in the number of peaks classified within a particular class and the total number of unique classes described. Non-flavonoid phenolics and flavonoids tended to be the most dominant chemical classes extracted across each genus, accounting for 50.5% and 17.9% of the total number of peaks described, respectively. Specifically, peaks were identified as alkaloids, fatty acids, flavonoids, non-flavonoid phenolics, and terpenoids in *Rosa*, *Magnolia,* and *Betula*; and alkaloids, fatty acids, non-flavonoid phenolics, and terpenoids in *Viburnum*.

**Figure 2.**
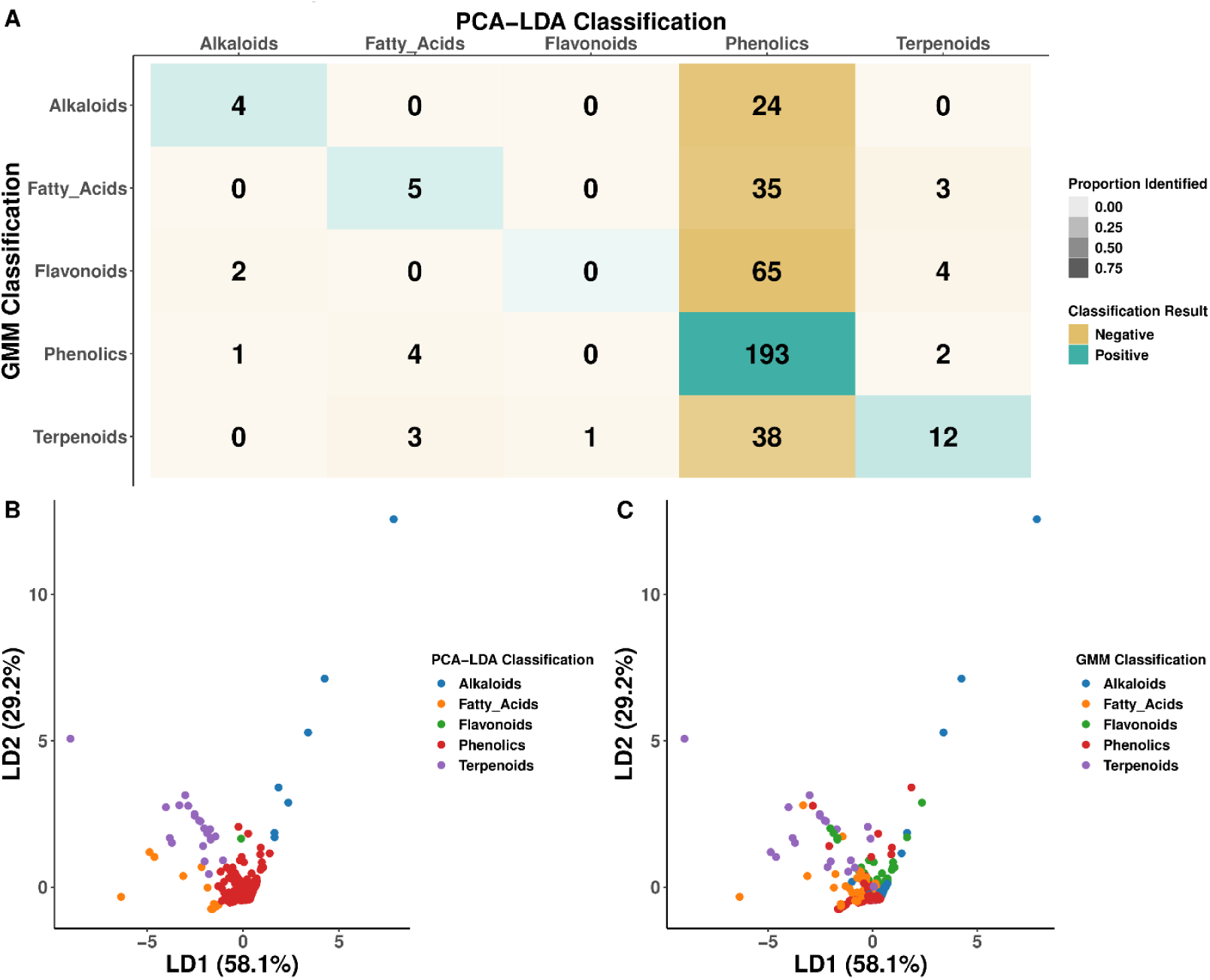
Model comparison between classification strength of a Gaussian Mixture Model (GMM) and Principal Component Analysis-Linear Discriminant Analysis (PCA-LDA). A) Confusion matrix comparing the number of predicted classifications made by the PCA-LDA model, against the accepted classifications made by the GMM. Opacity describes the proportion of compounds classified by the PCA-LDA relative to the classifications made by the GMM, with darker colors indicating greater proportions. Color indicates level of agreement (cyan being positive model agreement, and tan being negative model disagreement). B and C) Individual loading plots showing linear discriminant (LD) scores of each peak derived from LDA model, with points colored by the classifier described by the PCA-LDA model (B) and the GMM model (C). Axes are the first two linear discriminant axes describing 87.3% of the observed variation in the data set. Points represent individual peaks in the chromatogram.

Initially, PCA was used as a data reduction method. For *Magnolia*, PC1 and PC2 explained 58.7% and 22.7% of the total variance, respectively; for *Betula* 44.6% and 32.5%; for *Rosa* 62.7% and 27.1%; and for *Viburnum* 47.4% and 23.5%. Qualitative cluster classifications were assigned to each data point in the PCA plane, with ellipses drawn around the centroid of each GMM derived cluster. Cluster classifications were made by visual comparison with published absorbance spectra (Haghi & Hatami, 2010; Hisiger & Jolicoeur, 2007; Taniguchi et al., 2023a; Walker & Hawkins, 1952). Visually, GMM classification was able to differentiate between clusters that would otherwise have been poorly resolved through PCA dimension reduction alone (Figure 1; Appendix 1). Clusters were more easily discerned by PCA dimensions beyond the first and second principal components, with clearer separation between GMM-defined classes being observed when the third and fourth components were included (Appendix 1). Each genus contains its own unique level of chemical diversity: 57 unique peaks were described in the *Betula* sample set; 128 peaks in *Magnolia*; 165 peaks in *Rosa*; and 46 peaks in *Viburnum*.

PCA was also applied to the global data set to assess the ability of PCA for feature reduction in a subsequent LDA model. In the global analysis, while PC1 and PC2 together explained 82% of the variation, subsequent LDA revealed poor clustering ability. Despite LD1 and LD2 explaining together over 87% of the variation, the LDA model resulted in classifications with low agreement compared to those predicted by the GMM model (mean 54% agreement [95% CI 0.490-0.590], P-value = 0.087). Overall, the LDA model yielded classifications with low agreement compared to those generated by the GMM model, with only 54.04% agreement with the LDA prediction (P-value = 0.08735; 95% CI [0.48991, 0.590289]; Appendix 1). The PCA-LDA analysis resulted in a Cohen’s Kappa value of 0.141 and further supported a low level of agreement between GMM labeled classes and LDA predicted classes (Appendix 1). Specificity of the LDA model was high for alkaloids, fatty acids, flavonoids, and terpenoids (∼96%), but dropped to 17.3% for non-flavonoid phenolics. The LDA model was most sensitive to non-flavonoid phenolics (96.5%), but could not correctly identify any flavonoids, resulting in a sensitivity of 0% (Appendix 1).

PGLS regressions were performed to examine the effect that abundances of chemical classes have on the chemical diversity indices of each genus. In general, abundance and diversity (SDI) and dominance (ISI) were weakly associated with abundances of all chemical classes (Table 1). Fatty acids, however, consistently showed a strong effect across all diversity indices of the observed chemical profiles in both *Viburnum* and *Rosa* (Table 1). Within *Viburnum,* there is a strong positive effect (Beta ∼ -4 ±1.3, P < 0.001), and within *Rosa,* there is a strong negative effect (Beta ∼ 3±0.8, P < 0.01).

**Table 1.** Significant PGLS associations for diversity indices predicting chemical class abundances.

| Genus | Diversity Indices | Chemical Class | $\beta^5$ | SE<sup>6</sup> (±) | Sig<sup>7</sup> |
| --- | --- | --- | --- | --- | --- |
| *Betula* | JSI<sup>1</sup> | NFP<sup>4</sup> | 0.966 | 0.433 | * |
|  | SDI<sup>2</sup> | Terpenoids | 0.011 | 0.005 | * |
|  | ISI<sup>3</sup> | NFP | 0.031 | 0.012 | * |
|  |  | Terpenoids | 0.002 | 0.001 | * |
| *Magnolia* | SDI | Fatty Acids | -1.398 | 0.334 | *** |
|  |  | Terpenoids | -0.705 | 0.222 | ** |
|  | ISI | Alkaloids | 0.062 | 0.008 | *** |
|  |  | Fatty Acids | 0.146 | 0.035 | *** |
|  |  | NFP | 0.015 | 0.003 | *** |
|  |  | Terpenoids | 0.101 | 0.017 | *** |
| *Rosa* | JSI | Fatty Acids | -3.026 | 0.797 | *** |
|  | SDI | Alkaloids | 0.036 | 0.013 | ** |
|  |  | Fatty Acids | -0.720 | 0.053 | *** |
|  | ISI | Alkaloids | 0.004 | 0.001 | *** |
|  |  | Fatty Acids | -0.071 | 0.006 | *** |
| *Viburnum* | JSI | Alkaloids | 0.198 | 0.059 | ** |
|  |  | Fatty Acids | 4.538 | 1.288 | ** |
|  | SDI | Fatty Acids | -0.907 | 0.155 | *** |
|  | ISI | Fatty Acids | -0.109 | 0.022 | *** |

Notes: <sup>1</sup>JSI: Jaccard Similarity Index. <sup>2</sup>SDI: Shannon diversity Index. <sup>3</sup>ISI: Inverse Simpson Index. <sup>4</sup>NFP: non-flavonoid phenolics. <sup>5</sup> $\beta$ : Effect size derived by the model beta coefficient of the respective PGLS model. <sup>6</sup>Standard Error. <sup>7</sup>Sig: significance at \* = P < 0.05, \*\* = P < 0.01, \*\*\* = P < 0.001

## Discussion

The primary objective of this study was to utilize GMMs to provide a more comprehensive characterization of phytochemical profiles than would be possible with HPLC-DAD or HPLC-MS in the absence of authentic standards or high-quality database information on a specific compound. As plants produce hundreds of thousands of unique metabolites, class-level characterization and quantification of phytochemical diversity within a plant sample offers a level of information highly useful for addressing physiological, ecological, evolutionary, and taxonomic research questions. Our resulting method extracts such data based on reconstructed absorption spectra and places unknown peaks into one of five chemical classes: alkaloids, fatty acids, flavonoids, non-flavonoid phenolics, and terpenoids.

The GMM model was particularly good at describing peaks with similar absorbance features. Non-flavonoid phenolics and flavonoids, for example, share similar absorbance maxima at ∼355 nm, with the major difference detected in this analysis being the absence of a secondary feature in non-flavonoid phenolics at ∼280nm (Figure 2A, magenta and orange arrows, respectively).

Not surprisingly, it was practically impossible for an LDA model to discern between these two chromophores, resulting in only a single terpenoid being incorrectly described as a flavonoid. The LDA was also particularly biased towards phenolics (Figure 2A). Non-flavonoid phenolics are relatively abundant compounds and absorb light particularly well (González-González et al., 2019; Haghi & Hatami, 2010), and their occurrence could potentially overshadow less prominent features in the chromatogram due to coelution or matrix effects. In our analysis, non-flavonoid phenolics were the most common class of peak detected (n = 200); therefore, it is reasonable to assume that the LDA model may be biased towards classes with higher membership (Blagus et al., 2010).

Additionally, non-flavonoid phenolics appear to share similar absorbance features with terpenoids at 280nm, and with alkaloids at 355nm (Figure 1A). Although the singular value decomposition (SVD) process within PCA can be applied to low-variance data, the data reduction process may eliminate small differences in the absorbance spectra when considered as a whole. Thus, the discriminatory power of GMMs likely stems from the model’s ability to form clusters based on the variation within the entire spectral object. The nature of eigenvector extraction in the SVD step of a PCA is aimed at reducing high-dimensional data to its most important “components,” and those components can easily obscure the small variations in a full multiwavelength absorption spectrum that make it unique to any one compound class. Thus, we argue that the information retained by using GMM is vital to accurate discrimination between chemical compounds that share some prominent spectral features, as is the case with non-flavonoid phenolics and flavonoids (Figure 1A).

Overall, the three diversity metrics calculated showed a strong potential to describe the chemical diversity attributed to each genus, with GMM-derived chemical class descriptions adding interesting nuance to how the diversity of metabolites is distributed. *Rosa* showed significant negative correlations between the Jaccard, Shannon, and Inverse Simpson indices and the relative class abundance of fatty acids (Table 1), suggesting that fatty acid abundances tended to be more associated with highly diverse (SDI) profiles with relatively few dominant peaks (ISI) with most species across the genus exhibiting more similar chemical profiles (JSI). The Shannon and Inverse Simpson Indices also show very weak, significant positive associations with alkaloids (Table 1), suggesting that across the genus, alkaloids abundances tend to be associated with and more diverse profiles that are not dominated by singular peaks. However, extremely strong negative correlations were detected between alkaloids and all other chemical classes, potentially indicating a significant production tradeoff. Alkaloids are known to be expensive to produce (Shonle & Bergelson, 2000), and thus their synthesis may deplete the available pools of amino acids and lipids that would otherwise be used to produce compounds in other chemical classes, such as fatty acids and terpenoids (Rodrigues et al., 2023).

Deeper connections between chemical classes themselves can also be observed, where overall patterns across the pairwise chemical class correlations (i.e., the occurrence of alkaloids affecting the occurrence of fatty acids) reveal potential metabolic construction or regulatory links. For example, in *Magnolia*, significant positive correlations are observed between alkaloids, terpenoids, fatty acids, and non-flavonoid phenolic acids, revealing potential interdependency patterns among these classes (Appendix 1). Terpenoids, for example, serve as the backbones for many alkaloids as a component of terpene alkaloid biosynthesis (Eljounaidi et al., 2024), which could explain the significant effect observed (Beta = 1.664 ± 0.14, P-value < 0.001).

Classification ability is also limited by a model’s ability to differentiate between different chemical classes while leveraging available chemical abundances. While diode array detectors are particularly useful for measuring compounds with strong chromophores (Figure 1B), they can easily be biased by chromophores with disproportionately high absorbances in the chromatogram, either by obscuring peaks due to coelution (Sawikowska et al., 2021) or low abundances not providing a strong enough signal to be observed over the baseline noise (Ghaoui & Rothman, 1992). In other words, highly abundant peaks can increase the instrument’s limit of detection, rendering compounds known to be present in certain samples undetectable. For example, *Viburnum* is known to produce a wide variety of flavonoids, namely anthocyanins (Sharifi-Rad et al., 2021), but they were not detected. Similarly, in *Betula*, flavonoids were not detected despite many members of the genus being known to produce high concentrations of flavonoids (Isidorov et al., 2014). However, our GMM model was unable to describe any peaks belonging to the flavonoid class in either genus (Appendix 1). In a study on *Viburnum* leaves, fruits, and flowers, it was found that the highest concentration of flavonoids was found in the flowers and fruits, while chlorogenic acid and other non-flavonoid phenolics dominated the metabolite profile of the leaves (Goławska et al., 2023). Thus, it is very likely that the lack of representation of notable flavonoids is due to sample preparation during the oven-drying process (Snoussi et al., 2021), or due to only the leaves being studied in our analysis. Further optimization of the chromatographic method could enhance the ability to differentiate between flavonoids and phenolics, as well as improve tissue selection and preparation.

Further improvements to this method could be achieved through the addition of analytical standards, which would help to distinguish sub-classifications within the main chemical classes described in this analysis. For example, the flavonoids class encompasses a very broad range of chemical compounds, all of which are differentiated by a single functional group modification and the position of double bonds on the ring structure of the molecule, resulting in almost identical absorption spectra across subgroups (Taniguchi et al., 2023). Additionally, publicly available libraries of absorbance spectra, such as those created by the National Institute of Standards and Technology (Gaithersburg, MD) and the Mass Bank of North America (https://mona.fiehnlab.ucdavis.edu/), for untargeted mass spectrometry data analysis could further enhance the identification of these classes of molecules. One alternative to utilizing analytical standards could be to employ sub-clustering on the initially described chemical classes; however, accurate reference material will still need to be available for such structural distinctions to be made with confidence.

## Conclusions

HPLC-DAD technology provides a powerful tool for observing photoactive compounds commonly produced by plant species. We employed an untargeted approach to observe coarse-grain chemical class information through a Gaussian finite mixture model clustering, which extends the use of this platform beyond targeted quantification. This chemical class information can be used to describe associations or trade-offs in metabolite production across species and relate chemical class abundance to other descriptors of chemical diversity, such as the Jaccard, Shannon, and Inverse Simpson’s Indices. This method could be further improved by subsequent sub-clustering within the chemical classes to observe more finite structural differences between different types of chemical species within a class, such as anthocyanin and flavanol composition within the flavonoid class. However, methods such as these would require more accurate analytical standards to build a more relevant spectral library, enabling the accurate and routine identification of compounds as different chemical species. The method we have presented here represents a push towards improving the accessibility of metabolomics. Our model can accurately detect small differences in absorbance patterns that would otherwise be overlooked, thereby improving the ability of plant metabolomics research to quickly and effectively measure chemical diversity beyond abundance and the number of peaks. Such feature extraction can be leveraged to observe broad chemical differences in a variety of systems beyond evolutionary chemical dynamics, such as crop genotype differences or differences between plant ecotypes.

## Contributions

JTG, JAD, and CMM designed the experiment. JAD constructed the phylogenies. JTG designed and performed the extractions and chromatography, processed the data, wrote the manuscript, and generated figures. JAD and CMM edited and contributed to the manuscript.

## Acknowledgements

We would like to thank undergraduates Isabelle Boykin, Kaley Gillan, Jenna Mosely, and Augie Antis for their assistance in preparing samples for this project. This work was supported by the Arnold Arboretum of Harvard University through the Katharine H. Putnam Fellowship in Plant Science awarded to C.M.M., and by start-up funding provided to C.M.M. by the University of Central Florida and to J.A.D. by Louisiana State University. Research support was also provided to J.T.G. by the Future Scholars Fellowship at Louisiana State University.

