## Appendix 1 for "Untargeted Metabolomics of Plant Samples using HPLC-DAD and Gaussian Mixture Models"

Appendix 1.1 Description of Chromatography methods for each genus used in study

**Commented [CM1]:** Not sure if you're permitted to do this for Appendices - (call them all Appendix 1 when it's supplemental methods, figures, and tables). If I recall correctly, both APPS and AJB have made me change this sort of thing before and give each item its own number.

Appendix 1.2 List of accepted taxon names and corresponding identifiers used by the Arnold Arboretum at Harvard University.

Appendix 1.3 List of NCBI Taxonomy IDs with corresponding species name given by the Arnold Arboretum. In the case where an exact match was not found by name, accepted synonyms were used based on the Taxonomic Name Resolution Service provided by BIEN (https://tnrs.biendata.org/).

Appendix 1.4 Principal Component Analysis and Linear Discriminant Analysis assessing the accuracy of chemical classifications against identifiers made by GMMs for *Magnolia*. Points represent individual peaks detected in the chromatogram. A) Individual plot of individual PC1 and PC2 values for each peak. Colors indicate represented classes. Ellipses show 95% confidence interval describing overall group variability within each chemical class with n>3 individuals. B) Plot of individual scores corresponding to the first two Linear Discriminant axes. Axes comprise 92.7% of the observed variation in the data set. Colors are assigned based on the chemical class described by Gaussian Mixture Model clustering.

Appendix 1.5 Principal Component Analysis and Linear Discriminant Analysis assessing the accuracy of chemical classifications against identifiers made by GMMs for *Rosa*. Points represent individual peaks detected in the chromatogram. A) Individual plot of individual PC1 and PC2 values for each peak. Colors indicate represented classes. Ellipses show 95% confidence interval describing overall group variability within each chemical class with n>3 individuals. B) Plot of individual scores corresponding to the first two Linear Discriminant axes. Axes comprise 95.5% of the observed variation in the data set. Colors are assigned based on the chemical class described by Gaussian Mixture Model clustering.

Appendix 1.6 Principal Component Analysis and Linear Discriminant Analysis assessing the accuracy of chemical classifications against identifiers made by GMMs for *Betula*. Points represent individual peaks detected in the chromatogram. A) Individual plot of individual PC1 and PC2 values for each peak. Colors indicate represented classes. Ellipses show 95% confidence interval describing overall group variability within each chemical class with n>3 individuals. B) Plot of individual scores corresponding to the first two Linear Discriminant axes. Axes comprise 99.9% of the observed variation in the data set. Colors are assigned based on the chemical class described by Gaussian Mixture Model clustering.

Appendix 1.7 Principal Component Analysis and Linear Discriminant Analysis assessing the accuracy of chemical classifications against identifiers made by GMMs for Viburnum. Points represent individual peaks detected in the chromatogram. A) Individual plot of individual PC1 and PC2 values for each peak. Colors indicate represented classes. Ellipses show 95% confidence interval describing overall group variability within each chemical class with n>3 individuals. B) Plot of individual scores corresponding to the first two Linear Discriminant axes.

Axes comprise 99.9% of the observed variation in the data set. Colors are assigned based on the chemical class described by Gaussian Mixture Model clustering

Appendix 1.8 Scatterplot matrix of the first four principal components of the global identified chemical data set. Each point represents an individual peak of all samples in the chromatogram. Colors are assigned based on the chemical class described by Gaussian Mixture Model clustering.

Appendix 1.9 LDA Model Statistics depicting sensitivity, specificity, Cohen's Kappa, accuracy, and chemical class prediction quality.

Appendix 1.10 PGLS correlations of diversity metrics predicting chemical class abundances. Beta coefficients show the effect size of the abundance of each chemical class predicting the diversity metric.

Appendix 1.11 Pairwise PGLS coefficients of associations between abundances of chemical classes. Beta coefficients show the effect size of abundance of Chemical Class A predicting the abundance of Chemical Class B. Bolded p-values indicate significant associations ($p < 0.05$).

**Commented [CM2]:** More detail is needed. Each data point represents what? One peak? One sample? One species? Is this across all four genera? Each figure needs to largely stand alone apart from the main text, so please over-explain to ensure a reader can tell what the graph represents.

**Commented [CM3]:** Formatting - each of your Appendices needs a stand-alone legend. You have buried that into the table. Please revise to fit journal style.

**Commented [CM4]:** The variable here is not clearly stated. What is being correlated???

### Appendix 1.1 Descriptions of High Performance Liquid Chromatography parameters used for each genus

Solvent A was a mixture of 3:97 LCMS grade Water:Acetonitrile

Solvent B was a mixture of 3:97 Acetonitrile:LCMS grade Water

Solvents were purchased from VWR Scientific (Randor, PA).

*Betula*

Solvent gradients for *Betula* were as follows: an initial linear gradient from 0-10% solvent D for 3 minutes; linear gradient from 10-35% solvent D for 5 minutes; linear gradient from 35-100% solvent D for 3 minutes. This was followed by a solvent switch step for 4 minutes, with an isocratic hold at 100% solvent C for 1 min. HPLC column compartment was held at room temperature (~25°C). Flow rate was held at 2 mL/min. Injection volume was 5 uL. A diode array detector was set to collect data from 190-900 nm, with a wavelength step of 5 nm. The method was run on an Agilent ZORBAX XDB C-18 600bar column, with inner diameter of 4.6 mm, length of 50 mm, and pore size of 1.8 micrometers (Santa Clara, CA).

*Magnolia*

Solvent gradients for *Magnolia* were as follows: an initial linear gradient from 0-3% solvent D for 3 minutes; linear gradient from 3-10% solvent D for 1 minute; linear gradient from 10-20% solvent D for 8 minutes; linear gradient from 20-100% solvent D for 3 minutes; and a solvent switch step for 5 minutes. HPLC column compartment was held at room temperature (~25°C). Flow rate was held at 1 mL/min. Injection volume was 3 uL. A diode array detector was set to collect data from 190-900 nm, with a wavelength step of 5 nm. The method was run on an Agilent Poroshell 120 EC C-18 column, with inner diameter of 3 mm, length of 150 mm, and pore size of 2.7 micrometers (Santa Clara, CA).

*Rosa*

Solvent gradients for *Rosa* were as follows: an initial linear gradient from 0 to 3% solvent D over 2 minutes; linear gradient from 3-17% solvent D over 1 minute; linear gradient from 17-18% solvent D over 12 minutes; linear gradient from 18-45% solvent D for 2 minutes; linear gradient from 45-70% solvent D for 10 minutes; linear gradient from 70-100% solvent D for 3 minutes. This was followed by an isochratic cleaning step at 100% solvent D for 5 minutes, and a solvent switch for 5 minutes. HPLC column compartment was held at 45°C. Flow rate was held at 0.5 mL/min. Injection volume was 3 uL. A diode array detector was set to collect data from 190-900 nm with a wavelength step of 5 nm. The method was run on an Agilent Poroshell 120 EC C-18

**Commented [CM5]:** Generally a supplemental methods section like this would be in paragraph form - I don't see a need for the section headers here, rather what would work better would be an introductory paragraph incorporating the solvent information followed by the per-genus paragraphs without headers.

**Commented [DJA6R5]:** i like this style

**Commented [DJA7]:** unless solvent A & B are used change the names of C & D column, with inner diameter of 3 mm, length of 150 mm, and pore size of 2.7 micrometers (Santa Clara, CA).

*Viburnum*

Solvent gradients for *Viburnum* were as follows: an initial linear gradient from 0-5% solvent D for 1 minute; linear gradient from 5-20% solvent D for 5 minutes; linear gradient from 20-25% solvent D for 2 minutes; and a linear gradient from 25-100% solvent D for 2 minutes. This was followed by a solvent switch step for 3 minutes, with an isocratic hold at 100% solvent C for 1 min. HPLC column compartment was held at room temperature (~25°C). Flow rate was held at 2 mL/min. Injection volume was 5 uL. A diode array detector was set to collect data from 190-900 nm, with a wavelength step of 5 nm. The method was run on an Agilent ZORBAX XDB C-18 600bar column, with inner diameter of 4.6 mm, length of 50 mm, and pore size of 1.8 micrometers (Santa Clara, CA).

Appendix 1.2 List of accepted taxon names and corresponding identifiers used by the Arnold Arboretum at Harvard University.

| Taxon Name | Accession |
|---|---|
| *Betula grossa* | 3114*A |
| *Betula schmidtii* | 241-2001*B |
| *Betula apoiensis* | 68-92*D |
| *Betula papyrifera* | 573-2008*A |
| *Betula ermanii* | 1172-88*A |
| *Betula populifolia* | 595-2008*A |
| *Betula occidentalis* var. *occidentalis* | 1314-79_A |
| *Betula litwinowii* | 1374-80*A |
| *Betula occidentalis* var. *occidentalis* | 389-2007*A |
| *Betula apoiensis* | 1427-77*H |
| *Betula ermanii* var. *ermanii* | 37-2005*C |
| *Betula uber* | 1438-84*A |
| *Betula litwinowii* | 1374-80*D |
| *Betula dahurica* | 364-97*A |
| *Betula nigra* | 1251-79*A |
| *Betula papyrifera* var. *subcordata* | 311-88*A |
| *Betula schmidtii* | 1422-77*E |
| *Betula populifolia* | 193-89*A |
| *Betula alleghaniensis* | 299-85*B |
| *Betula schmidtii* | 251-98*B |
| *Betula grossa* | 199-2007*A |
| *Betula nigra* | 1199*N |
| *Betula pubescens* var. *tortuosa* | 410-84*B |
| *Betula platyphylla* var. *japonica* | 72-92*B |
| *Betula ablo-sinensis* | 121-2007*B |
| *Betula utilis var. utilis* | 355-2001*A |
| *Betula costata* | 5-95*A |
| *Betula populifolia* | 127-2005*B |
| *Betula dahurica* | 1015-80*A |
| *Betula pumila* | 800-93*A |
| *Betula pubescens* | 439-85*C |
| *Betula pendula* | 105-96*C |
| *Betula lenta* | 628-88*A |
| *Betula globispica* | 10-88*A |
| *Betula ablo-sinensis* | 121-2007*A |
| *Betula uber* | 1433-84*B |
| *Betula fruticosa* | 307-2008*A |
| *Betula litwinowii* | 1347-77*A |

**Commented [CM8]:** For appendices, use Page Breaks between each Appendix.

**Commented [CM9]:** Italicize scientific names (but not "var." or "ssp."). Also - why bold?

**Commented [JG10R9]:** Was just an aesthetics choice, but I resolved the italics issues

| | |
|---|---|
| *Betula nigra* | 1199*D |
| *Betula chinensis* | 4-95*A |
| *Betula ermanii* | 351-97*A |
| *Betula pendula* | 105-96*A |
| *Betula lenta* | 126-2005*A |
| *Betula alleghaniensis* | 104-79*B |
| *Betula occidentalis* var. *occidentalis* | 1314-79*C |
| *Betula whilis* var. *whilis* | 355-2001*B |
| *Betula pubescens* | 353-93*B |
| *Betula mandshurica* | 465-94*A |
| *Betula chichibuensis* | 389-93*A |
| *Betula dahurica* | 1438-77*A |
| *Betula jacquemontii* | 74-94*A |
| *Betula alleghaniensis* | 566-2008*B |
| *Betula maximowicziana* | 1078-76*A |
| *Betula maximowicziana* | 1118-76*A |
| *Betula lenta* | 17679*A |
| *Betula pumila* | 800-93*B |
| *Betula pendula* | 1319-85*A |
| *Betula costata* | 373-97*B |
| *Magnolia acuminata* | 71-88*C |
| *Magnolia sargentiana* | 155-96*A |
| *Magnolia sieboldii* | 404-97*B |
| *Magnolia macrophylla* ssp. *macrophylla* | 961-89*A |
| *Magnolia liliiflora* | 374-2001*B |
| *Magnolia acuminata subcordata* | 15157*D |
| *Magnolia macrophylla* ssp. *macrophylla* | 299-2001*A |
| *Magnolia zenii* | 549-2009*A |
| *Magnolia sieboldii* | 589-87*A |
| *Magnolia obovata* | 470-90*A |
| *Magnolia denudata* | 1165-84*B |
| *Magnolia sieboldii* | 404-97*C |
| *Magnolia officinalis biloba* | 398-81*F |
| *Magnolia macrophylla* ssp. *ashei* | 396-96*B |
| *Magnolia kobus* | 141-41*A |
| *Magnolia acuminata* | 137-99*A |
| *Magnolia denudata* | 344-81*A |
| *Magnolia macrophylla* ssp. *ashei* | 396-96*A |
| *Magnolia cylindrica* | 397-81*A |
| *Magnolia officinalis* | 158-96*A |
| *Magnolia salicifolia* | 392-2008*A |

| | |
|---|---|
| *Magnolia tripetala* | 561-2009*A |
| *Magnolia salicifolia* | 874-28*F |
| *Magnolia macrophylla* ssp. *macrophylla* | 960-89*A |
| *Magnolia virginiana australis* | 1275-80*A |
| *Magnolia sprengeri* | 228-2005*A |
| *Magnolia fraseri* | 23-88*C |
| *Magnolia fraseri* | 999-79*A |
| *Magnolia virginiana* | 762-83*B |
| *Magnolia virginiana* | 420-92*B |
| *Magnolia virginiana australis* | 1275-80*B |
| *Magnolia kobus* | 294-57*A |
| *Magnolia stellata* | 393-2008*A |
| *Magnolia zenii* | 430-91*A |
| *Magnolia amoena* | 56-96*A |
| *Magnolia stellata* | 803-86*A |
| *Magnolia acuminata* | 136-99*A |
| *Magnolia acuminata subcordata* | 732-90*C |
| *Magnolia officinalis biloba* | 160-96*A |
| *Magnolia macrophylla* | 560-2009*C |
| *Magnolia stellata* | 1046-78*A |
| *Magnolia kobus* | 1199-77*E |
| *Magnolia cylindrica* | 158-83*B |
| *Magnolia macrophylla* | 560-2009*A |
| *Magnolia cylindrica* | 1096-89*A |
| *Magnolia tripetala* | 562-2009*A |
| *Magnolia officinalis biloba* | 398-81*F |
| *Magnolia salicifolia* | 979-86*A |
| *Magnolia macrophylla* ssp. *ashei* | 1067-64*A |
| *Magnolia liliiflora* | 375-2001*A |
| *Magnolia officinalis* | 472-2010*A |
| *Magnolia macrophylla* | 560-2009*B |
| *Magnolia fraseri* | 964-79*C |
| *Magnolia virginiana* | 613-39*A |
| *Magnolia denudata* | 1165-84*D |
| *Magnolia acuminata subcordata* | 15157*A |
| *Magnolia tripetala* | 426-76*A |
| *Rosa roxbighii* | 253-83*MASS |
| *Rosa setisera* | 547-2009*A |
| *Rosa spinosissime* | 1102-83*MASS |
| *Rosa woodsii ultramontana* | 1195-82*MASS |
| *Rosa cinnamomca* | 383-79*MASS |

| | |
|---|---|
| *Rosa amblyotis* | 234-84*MASS |
| *Rosa blanda* | 688-82*MASS |
| *Rosa nutkana nutkana* | 184-2009*A |
| *Rosa henryi* | 117-98*A |
| *Rosa amblyotis* | 1521-83*MASS |
| *Rosa serafinii* | 700-82*MASS |
| *Rosa corymbitera* | 620-78*B |
| *Rosa bella* | 455-83*A |
| *Rosa sherardii* | 420-80*MASS |
| *Rosa gymnocarpa* | 1005-85*A |
| *Rosa gallica officinalis* | 270-94*B |
| *Rosa alberti* | 644-83*MASS |
| *Rosa omeiensis* | 63-96*A |
| *Rosa villosa* | 1305-80*MASS |
| *Rosa palustris* | 1179-83*MASS |
| *Rosa eglanteria* | 389-94*C |
| *Rosa canina* | 1085-85*A |
| *Rosa nitidula* | 348-94*A |
| *Rosa arvensis* | 579-94*A |
| *Rosa prattii* | 852-80*MASS |
| *Rosa hugonis* | 321-84*MASS |
| *Rosa palustris* | 163-85*MASS |
| *Rosa woodsii ultramontana* | 517-78*MASS |
| *Rosa macrocarpa* | 600-97*B |
| *Rosa dumalis dumalis* | 840-80*MASS |
| *Rosa iberica* | 728-89*A |
| *Rosa majalis* | 621-2007*A |
| *Rosa nutkana hispida* | 516-78*MASS |
| *Rosa moyesii* | 460-87*MASS |
| *Rosa stylosa* | 249-94*B |
| *Rosa acicularis* | 100-77*MASS |
| *Rosa rugosa* | 67-2002*A |
| *Rosa schrenkara* | 839-90*A |
| *Rosa pisocarpa* | 1760-81*MASS |
| *Rosa virginana* | 105-2006*MASS |
| *Rosa sweginozii* | 1109-88*B |
| *Rosa carolina* | 41-88*A |
| *Rosa foetida bicolor* | 77-2012*C |
| *Rosa mollis* | 61-2009*B |
| *Rosa zalana* | 928-78*G |
| *Rosa corymbitfera* | 388-94*A |

| | |
|---|---|
| *Rosa prattii* | 543-90*F |
| *Rosa canina* | 294-2008*C |
| *Rosa arkansana* | 314-90*MASS |
| *Rosa rugosa* | 72-90*MASS |
| *Rosa nitida* | 370-84*MASS |
| *Rosa scabriuscula* | 390-94*H |
| *Rosa nitida* | 1007-81*MASS |
| *Rosa pendulina* | 993-3*A |
| *Rosa primula* | 276-85*MASS |
| *Rosa nitida* | 483-97*MASS |
| *Rosa canina* | 387-94*C |
| *Rosa mollis* | 289-87*A |
| *Rosa woodii* | 266-90*A |
| *Rosa foetida bicolor* | 77-2012*B |
| *Rosa coriifolia* | 716-82*MASS |
| *Rosa californica* | 1388-83*MASS |
| *Rosa spinossissima ataica* | 3136-2*A |
| *Rosa arvensis* | 948-79*MASS |
| *Rosa corymbitela* | 388-94*B |
| *Rosa oxyodon* | 295-2008*B |
| *Rosa multiflora* forma *roseiflora* | 1060-83*B |
| *Rosa setigera* | 374-84*A |
| *Rosa palustris* | 645-83*MASS |
| *Rosa glauca* | 347-94*A |
| *Rosa oxyodon* | 295-2008*A |
| *Rosa luciae* | 1556-83*MASS |
| *Rosa inodora* | 330-85*A |
| *Rosa transorrisonensis* | 258-98*C |
| *Rosa acicularis nipponensis* | 707-82*MASS |
| *Rosa acicularis* ssp. *sayi* | 515-78*B |
| *Rosa micrantha* | 1414-83*A |
| *Rosa dauurica* | 324-97*MASS |
| *Rosa cetifolia* | 327-84*C |
| *Rosa inodora* | 476-78*MASS |
| *Rosa spinosissima* | 428-85*A |
| *Rosa virginana* | 106-2006*A |
| *Rosa acicularis bourgequi* | 1049-83*MASS |
| *Rosa inodora* | 330-85*B |
| *Rosa rugosa* | 277-84*MASS |
| *Rosa woodsii* | 109-78*MASS |
| *Rosa spinosissima* | 1462-80*MASS |

| | |
|---|---|
| *Viburnum recognitum* | 633-93*A |
| *Viburnum plicatum* | 933-4*A |
| *Viburnum erubescens* var. *gracilipes* | 798-85*A |
| *Viburnum hupehense* ssp. *hupehense* | 80-81*B |
| *Viburnum mollc forma leiophyllum* | 167-2000*A |
| *Viburnum glomeratum* | 650-2010*A |
| *Viburnum lentago* | 301-86*A |
| *Viburnum rhytidophyllum* | 45-81*B |
| *Viburnum rhytidophyllum* | 1386-82*A |
| *Viburnum dilatatum* | 821-85*A |
| *Viburnum opulus* | 352-78*A |
| *Viburnum rafinesquianum* | 227-2006*A |
| *Viburnum lobophyllum* | 1166-83*A |
| *Viburnum lobophyllum* | 206-2004*A |
| *Viburnum setigerum* | 305-2002*A |
| *Viburnum opulus* var. *calvescens* | 719-88*A |
| *Viburnum wrightii* var. *eglandulosum* | 825-63*A |
| *Viburnum dentatum* var. *pubescens* | 269-32*A |
| *Viburnum flavescens* | 1964-80*A |
| *Viburnum plicatum* | 18016-1*A |
| *Viburnum sieboldii* | 616-6*B |
| *Viburnum scabrellum* | 911-79*MASS |
| *Viburnum bitchiuense* | 1797-77*A |
| *Viburnum bitchiuense* | 1797-77*C |
| *Viburnum rufidulum* | 3943-1*D |
| *Viburnum betulifolium* | 255-2001*A |
| *Viburnum ichangense* var. *abro* | 259-2001*A |
| *Viburnum plicatum* var. *tementosum* | 503-84*A |
| *Viburnum melanocarpum* | 386-81*D |
| *Viburnum erosum* | 619-88*A |
| *Viburnum lobophyllum* | 1875-80*A |
| *Viburnum rafinesquianum* | 480-97*A |
| *Viburnum sargentii* | 379-97*A |
| *Viburnum sargentii* | 1897-80*B |
| *Viburnum sieboldii* | 398-62*D |
| *Viburnum betulifolium* | 255-2001*B |
| *Viburnum buddleitolium* | 7533*B |
| *Viburnum veitchii* | 457-94*A |
| *Viburnum hanceanum* | 360-95*A |
| *Viburnum rafinesquianum affine* | 18295*A |
| *Viburnum veitchii* | 232-2006*B |

| | |
|---|---|
| *Viburnum bitchiuense* | 2047-77*A |
| *Viburnum carlesii* | 892-61*A |
| *Viburnum setigerum* | 1635-80*A |
| *Viburnum dentatum venosum* | 268-85*D |
| *Viburnum cassinoides* | 109-79*A |
| *Viburnum mongolicum* | 254-2001*A |
| *Viburnum prunifolium* | 1189-85*A |
| *Viburnum bracteatum* | 1067-87*A |
| *Viburnum dilatatum* | 138-52*A |
| *Viburnum recognitum* | 19-60*A |
| *Viburnum lentago* | 306-90*A |
| *Viburnum hanceanum* | 360-95*B |
| *Viburnum dentatum* | 5070-1*A |
| *Viburnum cassinoides* | 1104-81*A |
| *Viburnum rutidulum* | 21418*A |
| *Viburnum lantana discolor* | 1294-83*A |
| *Viburnum dentatum venosum* | 18008*MASS |
| *Viburnum lentago* | 1715-81*A |
| *Viburnum dentatum indianense* | 101-38*A |
| *Viburnum carlesii* | 17981-2*A |
| *Viburnum molle* | 18294*A |
| *Viburnum glomeratum* | 650-2010*C |
| *Viburnum rhytidophyllum* | 57-81*A |
| *Viburnum dentatum* | 293-85*A |
| *Viburnum opulus calvescens* | 287-2002*A |
| *Viburnum trilobum* | 1097-60*A |
| *Viburnum ichangense* | 51-60*A |
| *Viburnum bracteatum* | 6119*A |
| *Viburnum betulifolium* | 1736-80*A |
| *Viburnum dentatum* | 8-2012*A |
| *Viburnum erosum* | 963-85*A |
| *Viburnum opulus* | 873-85*A |
| *Viburnum sieboldii* | 616-6*A |
| *Viburnum furcatum* | 17988*A |
| *Viburnum cassinoides* | 874-85*A |
| *Viburnum burejaeticum* | 375-97*A |
| *Viburnum sargentii* | 79-90*B |
| *Viburnum wrightii* | 1825-77*MASS |
| *Viburnum trilobum* | 361-2006*A |
| *Viburnum burejaeticum* | 375-97*C |
| *Viburnum furcatum* | 17988*B |

| | |
|---|---|
| *Viburnum nudum* | 565-2003*B |
| *Viburnum trilobum* | 361-2006*C |
| *Viburnum acerifolium* | 364-84*A |
| *Viburnum recognitum* | 1471-83*A |
| *Viburnum hanceanum* | 360-95*C |
| *Viburnum prunifolium* | 237-2006*A |
| *Viburnum rafinesquianum* var. *affine* | 4622-2*B |
| *Viburnum hupehense* ssp. *hupehense* | 1748-80*A |
| *Viburnum molle* forma *leiophyllum* | 4643-1*A |
| *Viburnum utile* | 638-94*A |
| *Viburnum erosum* | 512-83*A |
| *Viburnum schensianum* | 832-63*MASS |
| *Viburnum hupehense* ssp. *hupehense* | 1985-80*A |
| *Viburnum lantana* | 206-96*A |
| *Viburnum veitchii* | 101-81*A |
| *Viburnum acerifolium* | 364-84*D |
| *Viburnum opulus* | 46-2006*A |
| *Viburnum bracteatum* | 1068-87*A |
| *Viburnum dentatum* var. *pubescens* | 1046-81*A |
| *Viburnum dilatatum* | 1804-77*A |
| *Viburnum schensianum* | 744-88*B |

Appendix 1.3 . List of NCBI Taxonomy IDs with corresponding species name given by the Arnold Arboretum. In the case where an exact match was not found by name, accepted synonyms were used based on the Taxonomic Name Resolution Service provided by BIEN (https://tnrs.biendata.org/).

| ***Taxon Name*** | **NCBI Accession Code** |
|---|---|
| *Betula albo-sinensis* | Betula_utilis_subsp_albosinensis_210132 |
| *Betula alleghaniensis* | Betula_alleghaniensis_21017 |
| *Betula apoiensis* | Betula_apoiensis_312781 |
| *Betula chichibuensis* | Betula_chichibuensis_312783 |
| *Betula chinensis* | Betula_chinensis_312784 |
| *Betula costata* | Betula_costata_253225 |
| *Betula dahurica* | Betula_davurica_253224 |
| *Betula ermanii* | Betula_ermanii_216992 |
| *Betula ermanii* var. *ermanii* | Betula_ermanii_var_ermanii_2900535 |
| *Betula fruticosa* | Betula_fruticosa_216993 |
| *Betula globispica* | Betula_globispica_312787 |
| *Betula grossa* | Betula_grossa_312788 |
| *Betula jacquemontii* | Betula_utilis_subsp_jacquemontii_1685999 |
| *Betula lenta* | Betula_lenta_216994 |
| *Betula litwinowii* | Betula_pubescens_var_litwinowii_1685994 |
| *Betula mandshurica* | Betula_pendula_subsp_mandshurica_1689654 |
| *Betula maximowicziana* | Betula_maximowicziana_211500 |
| *Betula nigra* | Betula_nigra_3508 |
| *Betula occidentalis* var. *occidentalis* | Betula_occidentalis_223244 |
| *Betula papyrifera* | Betula_papyrifera_3507 |
| *Betula papyrifera* var. *subcordata* | Betula_papyrifera_var_commutata_1685990 |
| *Betula pendula* | Betula_pendula_3505 |
| *Betula platyphylla* var. *japonica* | Betula_platyphylla_var_japonica_216988 |
| *Betula populifolia* | Betula_populifolia_216989 |
| *Betula pubescens* ssp. *tortuosa* | Betula_pubescens_38787 |
| *Betula pumila* | Betula_pumila_284674 |
| *Betula schmidtii* | Betula_schmidtii_216986 |
| *Betula uber* | Betula_uber_253223 |
| *Betula utilis ssp. utilis* | Betula_utilis_var_prattii_1686001 |
| *Magnolia_acuminata* | Magnolia_acuminata_3404 |
| *Magnolia_amoena* | Magnolia_amoena_86724 |
| *Magnolia_cylindrica* | Magnolia_cylindrica_86728 |
| *Magnolia_denudata* | Magnolia_denudata_85856 |
| *Magnolia_fraseri* | Magnolia_fraseri_85857 |
| *Magnolia_kobus* | Magnolia_kobus_54732 |
| *Magnolia_liliiflora* | Magnolia_liliiflora_3403 |

**Commented [CM11]:** Page break before each new Appendix.

**Commented [CM12]:** Again, italics?

| | |
|---|---|
| *Magnolia_macrophylla* | Magnolia_macrophylla_3410 |
| *Magnolia_macrophylla* ssp. *ashei* | Magnolia_ashei_86777 |
| *Magnolia_macrophylla* ssp. *macrophylla* | Magnolia_macrophylla_subsp_macrophylla_86778 |
| *Magnolia_obovata* | Magnolia_obovata_349509 |
| *Magnolia_officinalis* | Magnolia_officinalis_85864 |
| *Magnolia_officinalis biloba* | Magnolia_officinalis_subsp_biloba_85865 |
| *Magnolia_salicifolia* | Magnolia_salicifolia_3411 |
| *Magnolia_sargentiana* | Magnolia_sargentiana_86736 |
| *Magnolia_sieboldii* | Magnolia_sieboldii_85868 |
| *Magnolia_sprengeri* | Magnolia_sprengeri_86737 |
| *Magnolia_stellata* | Magnolia_stellata_54733 |
| *Magnolia_tripetala* | Magnolia_tripetala_44926 |
| *Magnolia_virginiana* | Magnolia_virginiana_3412 |
| *Magnolia_zenii* | Magnolia_zenii_86739 |
| *Rosa acicularis* | Rosa_acicularis_117260 |
| *Rosa acicularis* var. *nipponensis* | Rosa_acicularis_var_nipponensis_283481 |
| *Rosa albertii* | Rosa_albertii_396727 |
| *Rosa amblyotis* | Rosa_amblyotis_1165904 |
| *Rosa arkansana* | Rosa_arkansana_267231 |
| *Rosa arvensis* | Rosa_arvensis_136998 |
| *Rosa bella* | Rosa_bella_396730 |
| *Rosa blanda* | Rosa_blanda_74640 |
| *Rosa californica* | Rosa_californica_74641 |
| *Rosa canina* | Rosa_canina_74635 |
| *Rosa carolina* | Rosa_carolina_74638 |
| *Rosa centifolia* | Rosa_x_centifolia_396733 |
| *Rosa coriifolia* | Rosa_caesia_267233 |
| *Rosa corymbifera* | Rosa_corymbifera_267235 |
| *Rosa davurica* | Rosa_davurica_237596 |
| *Rosa dumalis var. dumalis* | Rosa_dumalis_291166 |
| *Rosa eglanteria* | Rosa_rubiginosa_74636 |
| *Rosa foetida* var. *bicolor* | Rosa_foetida_74629 |
| *Rosa gallica* var. *officinalis* | Rosa_gallica_74632 |
| *Rosa glauca* | Rosa_glauca_74637 |
| *Rosa gymnocarpa* | Rosa_gymnocarpa_339615 |
| *Rosa henryi* | Rosa_henryi_117262 |
| *Rosa hugonis* | Rosa_hugonis_267238 |
| *Rosa inodora* | Rosa_elliptica_subsp_inodora_323239 |
| *Rosa macrocarpa* | Rosa_odorata_var_gigantea_74650 |
| *Rosa majalis* | Rosa_majalis_57935 |
| *Rosa micrantha* | Rosa_micrantha_267241 |

| | |
|---|---|
| *Rosa mollis* | Rosa_mollis_267242 |
| *Rosa moyesii* | Rosa_moyesii_74644 |
| *Rosa multiflora* forma *roseiflora* | Rosa_multiflora_74647 |
| *Rosa nitida* | Rosa_nitida_267245 |
| *Rosa nitidula* | Rosa_x_nitidula_1608484 |
| *Rosa nutkana* var. *hispida* | Rosa_nutkana_var_hispida_396746 |
| *Rosa nutkana* var. *nutkana* | Rosa_nutkana_var_nutkana_1608493 |
| *Rosa omeiensis* | Rosa_omeiensis_648851 |
| *Rosa palustris* | Rosa_palustris_267247 |
| *Rosa pendulina* | Rosa_pendulina_74642 |
| *Rosa pisocarpa* | Rosa_pisocarpa_339616 |
| *Rosa prattii* | Rosa_prattii_971288 |
| *Rosa primula* | Rosa_primula_267249 |
| *Rosa roxburghii* | Rosa_roxburghii_74654 |
| *Rosa rugosa* | Rosa_rugosa_74645 |
| *Rosa scabriuscula* | Rosa_tomentosa_267260 |
| *Rosa setigera* | Rosa_setigera_137000 |
| *Rosa sherardii* | Rosa_sherardii_267253 |
| *Rosa spinosissima* | Rosa_spinosissima_74630 |
| *Rosa spinosissima* var. *altaica* | Rosa_spinosissima_var_altaica_267230 |
| *Rosa stylosa* | Rosa_stylosa_267255 |
| *Rosa sweginzowii* | Rosa_sweginzowii_971290 |
| *Rosa transmorrisonensis* | Rosa_transmorrisonensis_1608497 |
| *Rosa villosa* | Rosa_villosa_267261 |
| *Rosa woodsii* | Rosa_woodsii_32246 |
| *Rosa woodsii* ssp. *ultramontana* | Rosa_woodsii_var_ultramontana_367856 |
| *Viburnum acerifolium* | Viburnum_acerifolium_4205 |
| *Viburnum betulifolium* | Viburnum_betulifolium_224736 |
| *Viburnum bitchiuense* | Viburnum_carlesii_var_bitchiuense_2819790 |
| *Viburnum bracteatum* | Viburnum_bracteatum_1127638 |
| *Viburnum buddleifolium* | Viburnum_buddleifolium_1190426 |
| *Viburnum burejaeticum* | Viburnum_burejaeticum_1190427 |
| *Viburnum carlesii* | Viburnum_carlesii_237927 |
| *Viburnum cassinoides* | Viburnum_cassinoides_1127640 |
| *Viburnum dentatum* | Viburnum_dentatum_61589 |
| *Viburnum dentatum* var. *indianense* | Viburnum_dentatum_var_indianense_1190429 |
| *Viburnum dentatum* var. *venosum* | Viburnum_dentatum_var_venosum_1190431 |
| *Viburnum dilatatum* | Viburnum_dilatatum_237933 |
| *Viburnum erosum* | Viburnum_erosum_237937 |
| *Viburnum erubescens* var. *gracilipes* | Viburnum_erubescens_237938 |
| *Viburnum flavescens* | Viburnum_flavescens_1127647 |

| | |
|---|---|
| *Viburnum furcatum* | Viburnum_furcatum_237940 |
| *Viburnum glomeratum* | Viburnum_glomeratum_1190434 |
| *Viburnum hanceanum* | Viburnum_hanceanum_1740872 |
| *Viburnum hupehense* ssp. *hupehense* | Viburnum_hupehense_1127649 |
| *Viburnum ichangense* | Viburnum_ichangense_1127650 |
| *Viburnum lantana* | Viburnum_lantana_237945 |
| *Viburnum lentago* | Viburnum_lentago_61590 |
| *Viburnum lobophyllum* | Viburnum_lobophyllum_237947 |
| *Viburnum melanocarpum* | Viburnum_melanocarpum_237948 |
| *Viburnum molle* | Viburnum_molle_237949 |
| *Viburnum mongolicum* | Viburnum_mongolicum_432677 |
| *Viburnum nudum* | Viburnum_nudum_237950 |
| *Viburnum opulus* | Viburnum_opulus_85293 |
| *Viburnum plicatum* | Viburnum_plicatum_179996 |
| *Viburnum plicatum* var. *tomentosum* | Viburnum_plicatum_var_tomentosum_237953 |
| *Viburnum prunifolium* | Viburnum_prunifolium_237954 |
| *Viburnum rafinesquianum* | Viburnum_rafinesqueanum_237955 |
| *Viburnum rafinesquianum* var. *affine* | Viburnum_rafinesqueanum_var_affine_1220048 |
| *Viburnum recognitum* | Viburnum_recognitum_1663608 |
| *Viburnum rhytidophyllum* | Viburnum_rhytidophyllum_47689 |
| *Viburnum rufidulum* | Viburnum_rufidulum_237956 |
| *Viburnum sargentii* | Viburnum_opulus_var_sargentii_237952 |
| *Viburnum scabrellum* | Viburnum_scabrellum_1190443 |
| *Viburnum schensianum* | Viburnum_schensianum_1127660 |
| *Viburnum setigerum* | Viburnum_setigerum_436501 |
| *Viburnum sieboldii* | Viburnum_sieboldii_61591 |
| *Viburnum trilobum* | Viburnum_opulus_var_americanum_237960 |
| *Viburnum utile* | Viburnum_utile_237963 |
| *Viburnum veitchii* | Viburnum_veitchii_1127665 |
| *Viburnum wrightii* | Viburnum_wrightii_349481 |

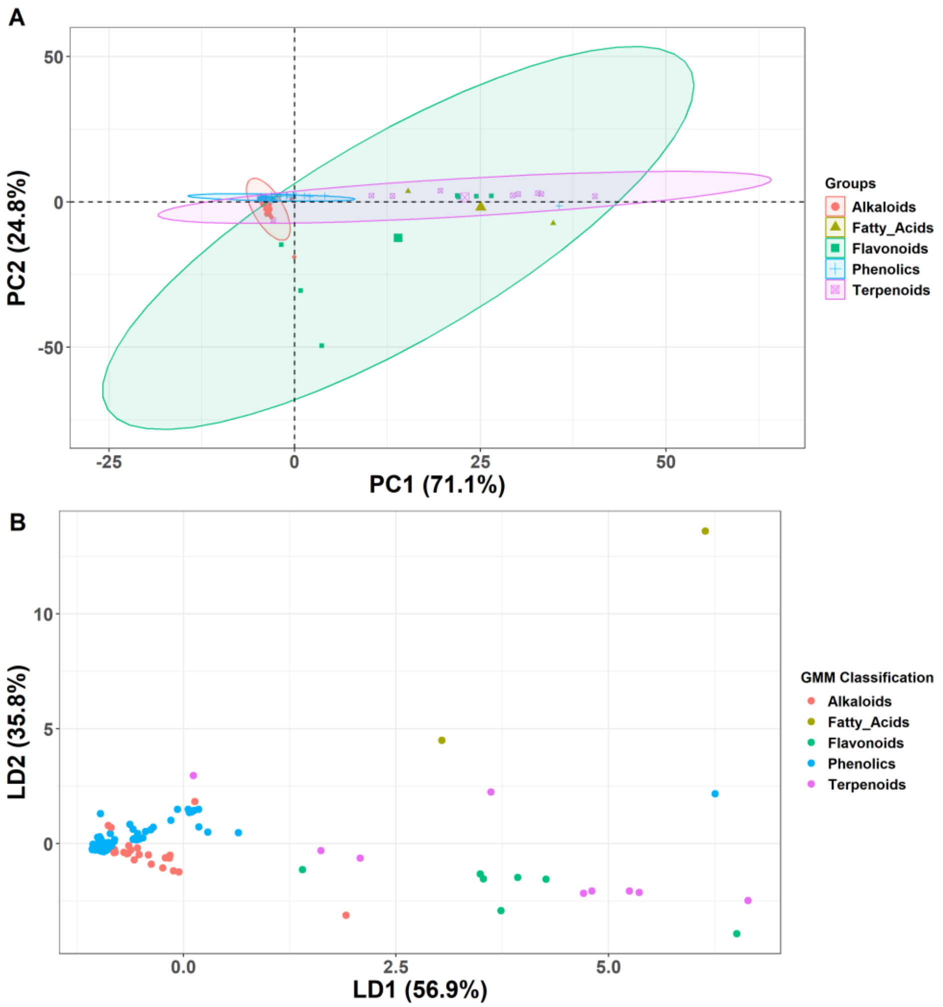


Appendix 1.4 Principal Component Analysis and Linear Discriminant Analysis assessing the accuracy of chemical classifications against identifiers made by Gaussian Mixture Model clusterings for *Magnolia*. Points represent individual peaks detected in the chromatogram. A) Individual plot of individual PC1 and PC2 values for each peak. Colors indicate represented classes. Ellipses represent 95% confidence intervals describing overall group variability within each chemical class, with n > 3 peaks per class. B) Plot of individual scores corresponding to the first two Linear Discriminant axes. Axes account for 92.7% of the observed variation in the dataset. Colors are assigned based on the chemical class described by the Gaussian Mixture Model clustering.

**Commented [DJA13]:** individuals or peaks?

**Commented [DJA14R13]:** peaks per class

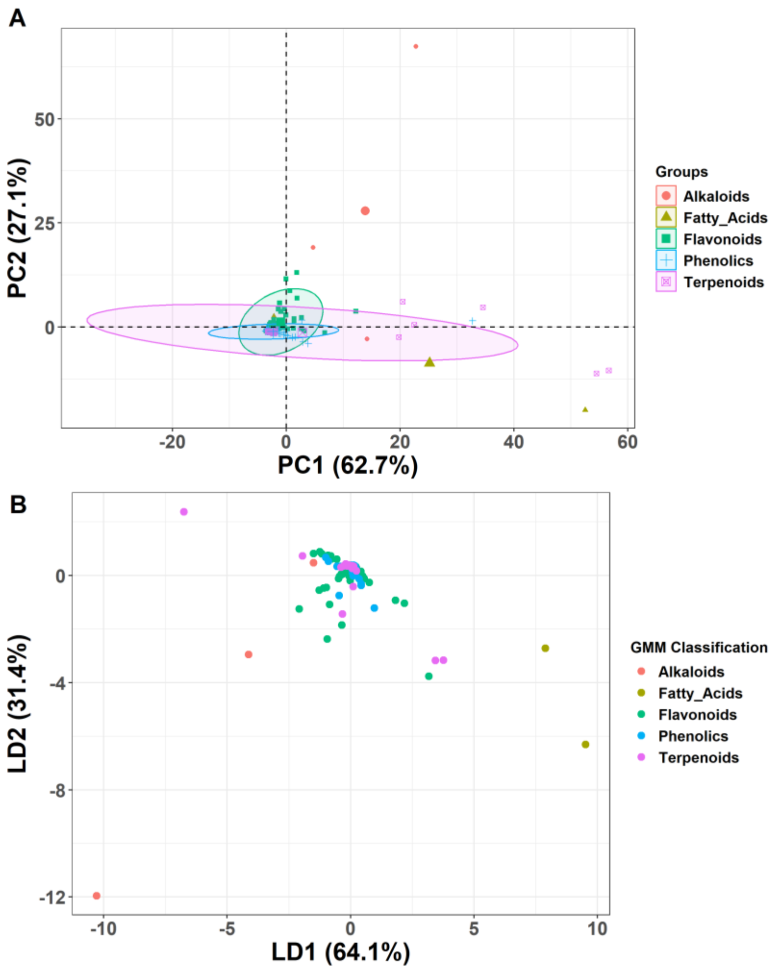


Appendix 1.5 Principal Component Analysis and Linear Discriminant Analysis assessing the accuracy of chemical classifications against identifiers made by Gaussian Mixture Model clusterings for *Rosa*. Points represent individual peaks detected in the chromatogram. A) Individual plot of individual PC1 and PC2 values for each peak. Colors indicate represented classes. Ellipses represent 95% confidence intervals describing overall group variability within each chemical class, with n > 3 peaks per class. B) Plot of individual scores corresponding to the first two Linear Discriminant axes. Axes comprise 95.5% of the observed variation in the data

set. Colors are assigned based on the chemical class described by the Gaussian Mixture Model clustering.

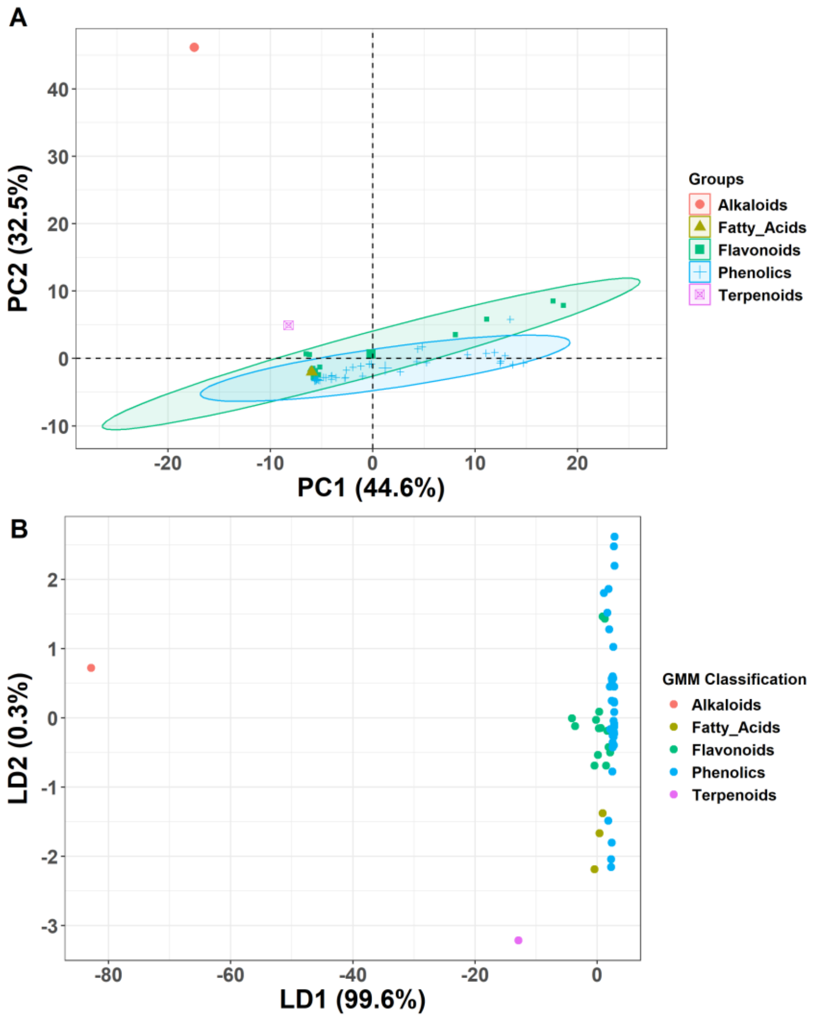


Appendix 1.6 Principal Component Analysis and Linear Discriminant Analysis assessing the accuracy of chemical classifications against identifiers made by Gaussian Mixture Model clusterings for *Betula*. Points represent individual peaks detected in the chromatogram. A) Individual plot of individual PC1 and PC2 values for each peak. Colors indicate represented classes. Ellipses represent 95% confidence intervals describing overall group variability within

each chemical class, with n > 3 peaks per class. B) Plot of individual scores corresponding to the first two Linear Discriminant axes. Axes comprise 99.9% of the observed variation in the data set. Colors are assigned based on the chemical class described by the Gaussian Mixture Model clustering.

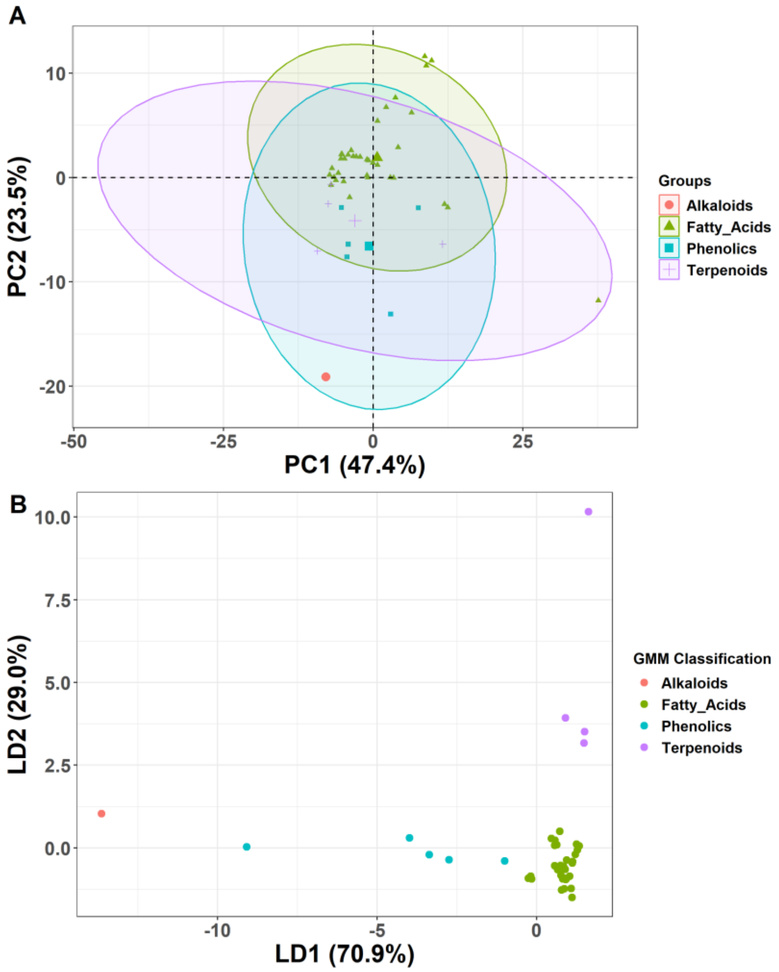


Appendix 1.7 Principal Component Analysis and Linear Discriminant Analysis assessing the accuracy of chemical classifications against identifiers made by Gaussian Mixture Model clusterings for *Viburnum*. Points represent individual peaks detected in the chromatogram. **A)** Individual plot of individual PC1 and PC2 values for each peak. Colors indicate represented classes. Ellipses represent 95% confidence intervals describing overall group variability within

**Commented [CM15]:** For all of your similarly formatted graphs, please improve the legends.

**Commented [CM16]:** So each point represents what? A peak, yes?

**Commented [JG17R16]:** Yeas each point is a different peak, added to the legend

**Commented [CM18]:** PC1 and PC2 values for each peak

each chemical class, with n > 3 individuals. **B)** Plot of individual scores corresponding to the first two Linear Discriminant axes. Axes comprise 99.9% of the observed variation in the data set. Colors are assigned based on the chemical class described by the Gaussian Mixture Model clustering.

**Commented [CM19]:** This is too brief/vague. It may feel redundant to you but you need to list everything relevant fro figure interpretation by a naïve reader. You're presenting the first two axes of linear discriminant analysis, which together explain 99% of the variation observed. Points again represent peaks.

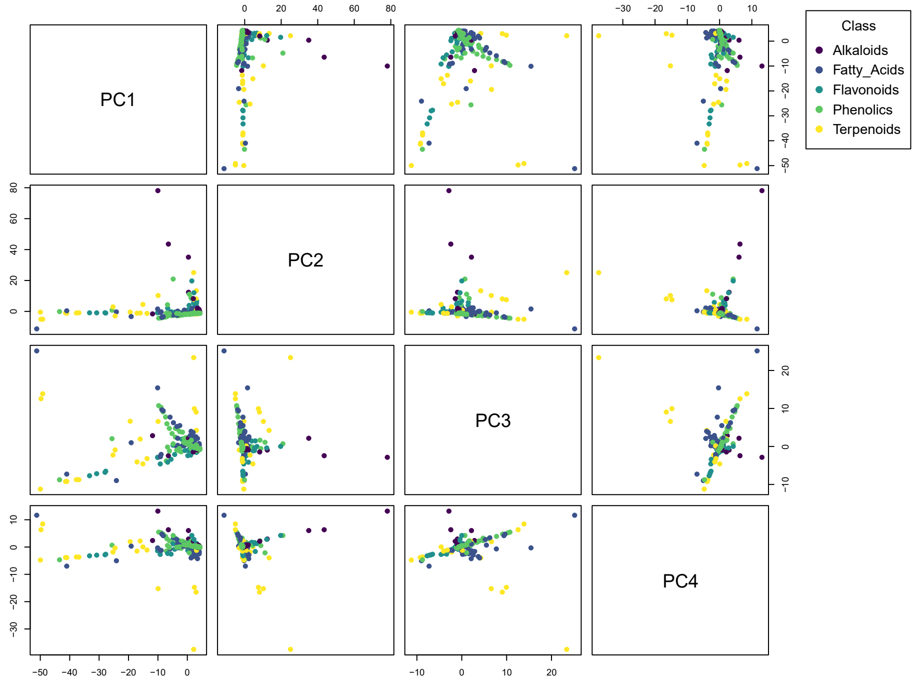


Appendix 1.8 Scatterplot matrix of the first four principal components of the global identified chemical data set. Each point represents an individual peak. Colors are assigned based on the chemical class described by Gaussian Mixture Model clustering.

**Commented [CM20]:** More detail is needed. Each data point represents what? One peak? One sample? One species? Is this across all four genera? Each figure needs to largely stand alone apart from the main text, so please over-explain to ensure a reader can tell what the graph represents.

Appendix 1.9 LDA Model Statistics depicting sensitivity, specificity, Cohen's Kappa, accuracy, and chemical class prediction quality.

**LDA model statistics, depicting sensitivity, specificity, Cohen's Kappa, and accuracy as measures of overall model performance. Confidence is described using the 95% confidence interval.**

| term | estimate | conf.low | conf.high | p.value |
|---|---|---|---|---|
| **accuracy** | 0.540404 | 0.489911 | 0.590289 | 0.08735 |
| **kappa** | 0.140967 | | | |

**LDA results of chemical class prediction quality, based on sensitivity, and specificity of identification using labels pre-described by GMM analysis.**

| | Alkaloids | Fatty Acids | Flavonoids | Phenolics | Terpenoids |
|---|---|---|---|---|---|
| Sensitivity | 0.142857 | 0.116279 | 0 | 0.965 | 0.222222 |
| Specificity | 0.991848 | 0.98017 | 0.996923 | 0.173469 | 0.973684 |
| Pos Pred Value | 0.571429 | 0.416667 | 0 | 0.543662 | 0.571429 |
| Neg Pred Value | 0.938303 | 0.901042 | 0.820253 | 0.829268 | 0.888 |
| Precision | 0.571429 | 0.416667 | 0 | 0.543662 | 0.571429 |
| Recall | 0.142857 | 0.116279 | 0 | 0.965 | 0.222222 |
| F1 | 0.228571 | 0.181818 | NA | 0.695495 | 0.32 |
| Prevalence | 0.070707 | 0.108586 | 0.179293 | 0.505051 | 0.136364 |
| Detection Rate | 0.010101 | 0.012626 | 0 | 0.487374 | 0.030303 |
| Detection Prevalence | 0.017677 | 0.030303 | 0.002525 | 0.896465 | 0.05303 |
| Balanced Accuracy | 0.567352 | 0.548225 | 0.498462 | 0.569235 | 0.597953 |
| Prevalence | 0.070707 | 0.108586 | 0.179293 | 0.505051 | 0.136364 |

**Commented [CM21]:** Formatting - each of your Appendices needs a stand-alone legend. You have buried that into the table. Please revise to fit journal style.

**Commented [JD22]:** what is the confidence percentile?

**Commented [JG23R22]:** 95 percent confidence interval

Appendix 1.10 PGLS correlations of diversity metrics predicting chemical class abundances. Beta coefficients show the effect size of the abundance of each chemical class predicting the diversity metric. Values are bolded if $P < 0.05$.

Commented [DJA24]: cant remember what your threshold was

| Genus | Diversity Metric | Class | Beta | Std. Error (±) | p-value |
|---|---|---|---|---|---|
| ***Betula*** | Jaccard Similarity Index | Alkaloids | 0.339 | 0.348 | 0.338964 |
| | | Flavonoids | 0.489 | 0.468 | 0.305506 |
| | | Phenolics | **0.966** | **0.433** | **0.034163** |
| | | Terpenoids | 0.042 | 0.026 | 0.122861 |
| | Shannon Index | Alkaloids | 0.099 | 0.062 | 0.12524 |
| | | Flavonoids | 0.022 | 0.088 | 0.802206 |
| | | Phenolics | 0.161 | 0.081 | 0.05802 |
| | | Terpenoids | **0.011** | **0.005** | **0.024535** |
| | Inverse Simpson Index | Alkaloids | 0.016 | 0.01 | 0.105356 |
| | | Flavonoids | 0.003 | 0.014 | 0.85312 |
| | | Phenolics | 0.031 | 0.012 | 0.015889 |
| | | Terpenoids | **0.002** | **0.001** | **0.020284** |
| ***Magnolia*** | Jaccard Similarity Index | Alkaloids | -0.51027 | 0.326199 | 0.134256 |
| | | Fatty Acids | -1.91791 | 0.937592 | 0.054897 |
| | | Flavonoids | 0.071661 | 0.077648 | 0.367634 |
| | | Phenolics | -0.02036 | 0.099632 | 0.840242 |
| | | Terpenoids | -1.02711 | 0.567224 | 0.086019 |
| | Shannon Index | Alkaloids | -0.24345 | 0.144573 | 0.108554 |
| | | Fatty Acids | **-1.39848** | **0.33405** | **0.000501** |
| | | Flavonoids | 0.054368 | 0.03323 | 0.118279 |
| | | Phenolics | 0.009857 | 0.044547 | 0.827234 |
| | | Terpenoids | **-0.70458** | **0.222084** | **0.005013** |
| | Inverse Simpson Index | Alkaloids | **0.061637** | **0.007983** | **2.83E-07** |
| | | Fatty Acids | **0.14571** | **0.035177** | **0.000554** |
| | | Flavonoids | -0.0008 | 0.003714 | 0.8326 |
| | | Phenolics | **0.014834** | **0.003203** | **0.000182** |
| | | Terpenoids | **0.101364** | **0.016954** | **9.40E-06** |
| ***Rosa*** | Jaccard Similarity Index | Alkaloids | 0.101746 | 0.106745 | 0.345002 |
| | | Fatty Acids | **-3.02619** | **0.796536** | **0.000388** |
| | | Flavonoids | -0.1024 | 0.223442 | 0.648707 |
| | | Phenolics | -0.16024 | 0.120796 | 0.190579 |
| | | Terpenoids | 0.181872 | 0.14403 | 0.212427 |
| | Shannon Index | Alkaloids | **0.035907** | **0.012623** | **0.006388** |
| | | Fatty Acids | **-0.71989** | **0.052892** | **1.31E-18** |

| | | | | | |
|---|---|---|---|---|---|
| | | Flavonoids | -0.00477 | 0.028243 | 0.866449 |
| | | Phenolics | -0.01273 | 0.015399 | 0.412358 |
| | | Terpenoids | -0.00199 | 0.018453 | 0.914335 |
| | Inverse Simpson Index | Alkaloids | **0.004414** | **0.001239** | **0.000806** |
| | | Fatty Acids | **-0.0706** | **0.006066** | **5.69E-16** |
| | | Flavonoids | -0.00016 | 0.002878 | 0.957121 |
| | | Phenolics | -0.00105 | 0.001573 | 0.508481 |
| | | Terpenoids | -0.00088 | 0.001876 | 0.641615 |
| ***Viburnum*** | Jaccard Similarity Index | Alkaloids | **0.197677** | **0.059273** | **0.001764** |
| | | Fatty Acids | **4.538316** | **1.287535** | **0.00102** |
| | | Phenolics | -0.00275 | 0.220467 | 0.990098 |
| | | Terpenoids | -0.22844 | 0.562047 | 0.68643 |
| | Shannon Index | Alkaloids | -0.00742 | 0.009366 | 0.432744 |
| | | Fatty Acids | **-0.907** | **0.154525** | **5.66E-07** |
| | | Phenolics | 0.042642 | 0.030595 | 0.170563 |
| | | Terpenoids | -0.06165 | 0.079338 | 0.441347 |
| | Inverse Simpson Index | Alkaloids | -0.0017 | 0.001217 | 0.170483 |
| | | Fatty Acids | **-0.10891** | **0.021725** | **9.70E-06** |
| | | Phenolics | 0.007677 | 0.003955 | 0.058811 |
| | | Terpenoids | -0.00333 | 0.010522 | 0.75335 |

Appendix 1.11 Pairwise PGLS coefficients of associations between abundances of chemical classes. Beta coefficients show the effect size of the abundance of Chemical Class A (explanatory) predicting the abundance of Chemical Class B (predictor). Bolded p-values indicate significant associations ($p < 0.05$).

Commented [CM25]: The variable here is not clearly stated. What is being correlated???

| Genus | Explanatory | Predictor | Beta | Std. Error (±) | t-value | p-value |
|---|---|---|---|---|---|---|
| **Betula** | Alkaloids | Flavonoids | 0.234 | 0.136 | 1.7276 | 0.0954794 |
| | Alkaloids | Non-Flavonoid Phenolics | 0.069 | 0.144 | 0.4777 | 0.636677 |
| | Alkaloids | Terpenoids | 1.241 | 2.450 | 0.5064 | 0.61666 |
| | Flavonoids | Alkaloids | 0.425 | 0.246 | 1.7276 | 0.0954794 |
| | Flavonoids | Non-Flavonoid Phenolics | 0.276 | 0.187 | 1.4735 | 0.152186 |
| | Flavonoids | Terpenoids | 4.730 | 3.190 | 1.4829 | 0.1496797 |
| | Non-Flavonoid Phenolics | Alkaloids | 0.122 | 0.255 | 0.4777 | 0.636677 |
| | Non-Flavonoid Phenolics | Flavonoids | 0.269 | 0.183 | 1.4735 | 0.152186 |
| | Non-Flavonoid Phenolics | Terpenoids | 3.796 | 3.194 | 1.1887 | 0.2449015 |
| | Terpenoids | Alkaloids | 0.008 | 0.015 | 0.5064 | 0.61666 |
| | Terpenoids | Flavonoids | 0.016 | 0.011 | 1.4829 | 0.1496797 |
| | Terpenoids | Non-Flavonoid Phenolics | 0.013 | 0.011 | 1.1887 | 0.2449015 |
| **Magnolia** | Alkaloids | Fatty Acids | 0.302 | 0.033 | 9.1927 | **2.01E-08** |
| | Alkaloids | Flavonoids | 0.360 | 0.998 | 0.3604 | 0.7224902 |
| | Alkaloids | Non-Flavonoid Phenolics | 2.609 | 0.526 | 4.9569 | **8.75E-05** |
| | Alkaloids | Terpenoids | 0.530 | 0.045 | 11.8815 | **3.06E-10** |
| | Fatty Acids | Alkaloids | 2.700 | 0.294 | 9.1927 | **2.01E-08** |
| | Fatty Acids | Flavonoids | -0.731 | 2.989 | -0.2444 | 0.8095356 |
| | Fatty Acids | Non-Flavonoid Phenolics | 6.331 | 1.888 | 3.3526 | **0.0033442** |
| | Fatty Acids | Terpenoids | 1.649 | 0.081 | 20.4702 | **2.09E-14** |
| | Flavonoids | Alkaloids | 0.019 | 0.052 | 0.3604 | 0.7224902 |
| | Flavonoids | Fatty Acids | -0.004 | 0.018 | -0.2444 | 0.8095356 |
| | Flavonoids | Non-Flavonoid Phenolics | 0.214 | 0.176 | 1.2185 | 0.2379618 |
| | Flavonoids | Terpenoids | -0.009 | 0.030 | -0.3120 | 0.7584629 |
| | Non-Flavonoid Phenolics | Alkaloids | 0.216 | 0.044 | 4.9569 | **8.75E-05** |
| | Non-Flavonoid Phenolics | Fatty Acids | 0.059 | 0.018 | 3.3526 | **0.0033442** |
| | Non-Flavonoid Phenolics | Flavonoids | 0.338 | 0.278 | 1.2185 | 0.2379618 |
| | Non-Flavonoid Phenolics | Terpenoids | 0.111 | 0.027 | 4.1129 | **0.0005921** |
| | Terpenoids | Alkaloids | 1.664 | 0.140 | 11.8815 | **3.06E-10** |
| | Terpenoids | Fatty Acids | 0.580 | 0.028 | 20.4702 | **2.09E-14** |
| | Terpenoids | Flavonoids | -0.552 | 1.771 | -0.3120 | 0.7584629 |
| | Terpenoids | Non-Flavonoid Phenolics | 4.226 | 1.028 | 4.1129 | **0.0005921** |
| **Rosa** | Alkaloids | Fatty Acids | -0.061 | 0.014 | -4.2847 | **8.11E-05** |

Commented [CM26]: If you are mixing standard notation and scientific notation, I recommend that you italicize or bold the significant ones.

| | | | | | | |
|---|---|---|---|---|---|---|
| | Alkaloids | Flavonoids | -0.208 | 0.061 | -3.4174 | **0.0012499** |
| | Alkaloids | Non-Flavonoid Phenolics | -0.418 | 0.108 | -3.8796 | **0.0003012** |
| | Alkaloids | Terpenoids | -0.204 | 0.099 | -2.0541 | **0.0451126** |
| | Fatty Acids | Alkaloids | -4.310 | 1.006 | -4.2847 | **8.11E-05** |
| | Fatty Acids | Flavonoids | 1.331 | 0.533 | 2.4994 | **0.0156994** |
| | Fatty Acids | Non-Flavonoid Phenolics | 2.751 | 0.953 | 2.8854 | **0.0057168** |
| | Fatty Acids | Terpenoids | 0.981 | 0.853 | 1.1499 | 0.2555625 |
| | Flavonoids | Alkaloids | -0.897 | 0.263 | -3.4174 | **0.0012499** |
| | Flavonoids | Fatty Acids | 0.082 | 0.033 | 2.4994 | **0.0156994** |
| | Flavonoids | Non-Flavonoid Phenolics | 0.928 | 0.220 | 4.2285 | **9.76E-05** |
| | Flavonoids | Terpenoids | -0.005 | 0.214 | -0.0226 | 0.9820392 |
| | Non-Flavonoid Phenolics | Alkaloids | -0.545 | 0.140 | -3.8796 | **0.0003012** |
| | Non-Flavonoid Phenolics | Fatty Acids | 0.051 | 0.018 | 2.8854 | **0.0057168** |
| | Non-Flavonoid Phenolics | Flavonoids | 0.280 | 0.066 | 4.2285 | **9.76E-05** |
| | Non-Flavonoid Phenolics | Terpenoids | 0.091 | 0.117 | 0.7812 | 0.4382899 |
| | Terpenoids | Alkaloids | -0.375 | 0.183 | -2.0541 | **0.0451126** |
| | Terpenoids | Fatty Acids | 0.026 | 0.022 | 1.1499 | 0.2555625 |
| | Terpenoids | Flavonoids | -0.002 | 0.091 | -0.0226 | 0.9820392 |
| | Terpenoids | Non-Flavonoid Phenolics | 0.129 | 0.166 | 0.7812 | 0.4382899 |
| **Viburnum** | Alkaloids | Fatty Acids | 0.013 | 0.007 | 1.9695 | 0.0553606 |
| | Alkaloids | Non-Flavonoid Phenolics | 0.030 | 0.046 | 0.6451 | 0.5222998 |
| | Alkaloids | Terpenoids | 0.035 | 0.017 | 2.0510 | **0.0463887** |
| | Fatty Acids | Alkaloids | 6.323 | 3.211 | 1.9695 | 0.0553606 |
| | Fatty Acids | Terpenoids | -0.159 | 0.395 | -0.4033 | 0.6887616 |
| | Fatty Acids | Non-Flavonoid Phenolics | 0.943 | 1.001 | 0.9420 | 0.351442 |
| | Non-Flavonoid Phenolics | Alkaloids | 0.325 | 0.503 | 0.6451 | 0.5222998 |
| | Non-Flavonoid Phenolics | Fatty Acids | 0.021 | 0.023 | 0.9420 | 0.351442 |
| | Non-Flavonoid Phenolics | Terpenoids | 0.185 | 0.053 | 3.5239 | **0.0010228** |
| | Terpenoids | Alkaloids | 2.528 | 1.233 | 2.0510 | **0.0463887** |
| | Terpenoids | Fatty Acids | -0.024 | 0.059 | -0.4033 | 0.6887616 |
| | Terpenoids | Non-Flavonoid Phenolics | 1.209 | 0.343 | 3.5239 | **0.0010228** |
